# Disrupting hierarchical control of nitrogen fixation enables carbon-dependent regulation of ammonia excretion in soil diazotrophs

**DOI:** 10.1101/2021.03.25.436926

**Authors:** Marcelo Bueno Batista, Paul Brett, Corinne Appia-Ayme, Yi-Ping Wang, Ray Dixon

## Abstract

The energetic requirements for biological nitrogen fixation necessitate stringent regulation of this process in response to diverse environmental constraints. To ensure that the nitrogen fixation machinery is expressed only under appropriate physiological conditions, the dedicated NifL-NifA regulatory system, prevalent in Proteobacteria, plays a crucial role in integrating signals of the oxygen, carbon and nitrogen status to control transcription of nitrogen fixation (*nif*) genes. Greater understanding of the intricate molecular mechanisms driving transcriptional control of *nif* genes may provide a blueprint for engineering diazotrophs that associate with cereals. In this study, we investigated the properties of a single amino acid substitution in NifA, (NifA-E356K) which disrupts the hierarchy of *nif* regulation in response to carbon and nitrogen status in *Azotobacter vinelandii*. The NifA-E356K substitution enabled overexpression of nitrogenase in the presence of excess fixed nitrogen and release of ammonia outside the cell. However, both of these properties were conditional upon the nature of the carbon source. Our studies reveal that the uncoupling of nitrogen fixation from its assimilation is likely to result from feedback regulation of glutamine synthetase, allowing surplus fixed nitrogen to be excreted. Reciprocal substitutions in NifA from other Proteobacteria yielded similar properties to the *A. vinelandii* counterpart, suggesting that this variant protein may facilitate engineering of carbon source-dependent ammonia excretion amongst diverse members of this family.

**Significance:** The NifL-NifA regulatory system provides dedicated signal transduction machinery to regulate nitrogen fixation in diverse Proteobacteria. Understanding how the balance of nitrogen and carbon resources is signalled via NifL-NifA for precise control of nitrogen fixation may lead to broadly applicable translational outputs. Here, we characterize a NifA variant that bypasses nitrogen regulation but is still dependent on the carbon status to enable ammonia excretion in soil diazotrophs. Disruption of the regulatory hierarchy in response to nitrogen and carbon suggests how the integration of environmental stimuli could be harnessed to engineer conditional release of fixed nitrogen for the benefit of cereal crops.

## Introduction

Biological nitrogen fixation requires diversion of reducing equivalents and ATP derived from carbon metabolism to support the high energetic demands of the enzyme nitrogenase, which converts dinitrogen to ammonia. Tight regulatory control provides the means to balance energy metabolism with nitrogen fixation and ammonia assimilation so that fixed nitrogen is not limiting under diazotrophic growth conditions. Achieving an appropriate balance between carbon and nitrogen metabolism is particularly important for diazotrophic bacteria in order to meet the energetic cost of nitrogen fixation, while thriving in competitive environments. While many studies on the regulation of nitrogen fixation have focused on intricate signalling mechanisms responding to the presence of oxygen and fixed nitrogen (1–3), regulation in response to the carbon status has not been extensively studied, despite its significance for the energetics of diazotrophy and the interplay required to balance the carbon : nitrogen ratio.

The nitrogen fixation specific, NifL-NifA, regulatory system provides a very sophisticated signal transduction complex for integration and transmission of various environmental cues to the transcriptional apparatus in *Azotobacter vinelandii* to regulate biosynthesis and expression of nitrogenase (reviewed in 8, 9). The anti-activator NifL is a multidomain protein carrying an N-terminal PAS domain (PAS1) that senses the redox status via a FAD co-factor (6, 7). A second PAS domain (PAS2) appears to play a structural role in relaying the redox changes perceived by the PAS1 domain to the central (H) and C-terminal (GHKL) domains of NifL (8, 9). The latter is responsible for ADP binding (10, 11) and is probably the site of interaction for the GlnK signal transduction protein, allowing integration of the nitrogen input into NifL-NifA regulation (12, 13). The protein partner of NifL, the prokaryotic enhancer binding protein NifA, which activates *nif* transcription, is comprised of an N-terminal regulatory domain (GAF), a central AAA+ sigma-54 activation domain and a C-terminal DNA binding domain. The regulatory GAF domain of *A. vinelandii* NifA binds 2-oxoglutarate (14), a TCA cycle intermediate at the interface of carbon and nitrogen metabolism (15). NifA can only escape inhibition by NifL, when the GAF domain is saturated with 2-oxoglutarate, thus potentially providing a mechanism for the NifL-NifA system to respond to the carbon status.

Understanding how the NifL-NifA system integrates diverse regulatory inputs may allow new strategies for engineering diazotrophs with enhanced ability to fix nitrogen and release ammonia to benefit crop nutrition. Ammonia excretion can be achieved in *A. vinelandii* by engineering constitutive expression of genes required for nitrogenase biosynthesis through inactivation of *nifL* (16–19). However, in the absence of active NifL, all the regulatory signal inputs that control NifA activity are removed, and as consequence, the resulting bacterial strain may be severely disadvantaged in the environment and even unstable under laboratory conditions as already reported (17, 19). Isolation of insertion mutants in *nifL* appears to be conditional upon second site mutations that may alter the level of *nif* gene expression. One such suppressor was identified in a promoter-like sequence upstream of *nifA*, presumably leading to a permissive reduction in *nifA* transcript levels (19). A full deletion of *nifL* has also been reported (18), but it is unclear if second site mutations occurred during its isolation.

One approach to potentially minimize the energetic burden associated with constitutive expression of nitrogenase is to ensure that regulatory control of NifA activity is maintained in energy-limiting environments. Random mutagenesis of *nifA* followed by screening the activity of the *A. vinelandii* NifL-NifA system in *E. coli* identified various NifA variants able to escape regulation by NifL under nitrogen excess conditions (20). One of these mutations, resulting in a charge-change substitution, E356K, located in the central catalytic domain of NifA, (hereafter named NifA-E356K), was found to require binding of 2-oxoglutarate to the GAF domain to escape NifL repression in response to excess fixed nitrogen (14, 21).

In this study we demonstrate that both expression and activity of nitrogenase are insensitive to the nitrogen status when the *nifA-E356K* mutation is introduced into *A. vinelandii*, resulting in excretion of ammonia at millimolar levels during exponential growth, a phenomenon correlated with feedback regulation of glutamine synthetase when nitrogenase is constitutively active. However, unregulated expression of *nif* genes and ammonia excretion by the *nifA-E356K* mutant is conditional on the nature of the carbon source indicating dependency on carbon status signalling and supporting previous biochemical observations that this NifA-E356K variant is dependent on the levels of 2-oxoglutarate to escape nitrogen regulation by NifL *in vitro* (21). Finally, we demonstrate that reciprocal substitutions in NifA proteins of other Proteobacteria lead to similar regulatory phenotypes when assayed in *E. coli* as a chassis and also when the substitution is engineered in the endophytic diazotroph *Pseudomonas stutzeri* A1501 (22, 23). In principle, this single amino acid substitution in NifA (E356K) provides a regulatory switch capable of activating *nif* gene expression under nitrogen excess conditions only when certain carbon sources are available in the environment, thus setting the foundation for engineering a synthetic symbiosis to deliver fixed nitrogen to cereal crops.

## Results

### The activity of NifA-E356K is not regulated in response to the nitrogen status in *A. vinelandii*, resulting in ammonia excretion

Previously the NifA-E356K variant protein was characterized either *in vivo* using *E. coli* as a chassis, or *in vitro* using purified protein components (20, 21). To evaluate if this variant protein would bypass NifL regulation in the original *A. vinelandii* DJ background, we introduced the *nifA-E356K* mutation into the chromosome and examined its influence on transcriptional regulation of the nitrogenase structural genes using RT-qPCR. As anticipated, no *nifH* transcripts were detected in the wild type strain (DJ) in the presence of either 25 or 5 mM of ammonium acetate, whilst increased transcript levels were observed in the absence of ammonium (Fig. 1A). In contrast, high levels of *nifH* transcripts were observed in all conditions tested for the *nifA-E356K* mutant strain (EK) (Fig. 1B), confirming that *nifA-E356K* is able to escape nitrogen regulation in *A. vinelandii*. Comparison of the levels of *nifH*, *nifL* and *nifA* transcripts in the EK strain relative to the wild type (DJ) (Fig. 1C) revealed that whilst *nifH* levels are higher in the EK mutant, this is not correlated with increased levels of *nifL* and *nifA* transcripts. This is in line with previous reports establishing that *nifLA* expression is not subject to autoactivation by NifA in *A. vinelandii* (24, 25) and suggests that constitutive expression of the nitrogenase structural gene operon in the EK mutant is intrinsic to the *nifA-E356K* mutation itself, rather than a consequence of overexpression of this mutant gene. The increase in *nifH* transcripts correlated with higher nitrogenase activity in the EK strain, which was not regulated in response to excess fixed nitrogen, in contrast to the wild-type strain (Fig. 1D). As anticipated from the energetic constraints associated with constitutive expression and activity of nitrogenase, the EK mutant had an apparent growth deficiency in liquid media (Supplementary Material Fig. S1). However, when an insertion replacing the *nifH* structural gene was introduced into the EK strain to generate strain EKH (EK, *ΔnifH::tetA)* this growth penalty was alleviated (Fig. S1-D-F) suggesting that it results from unregulated expression and activity of nitrogenase.

**Fig. 1.**
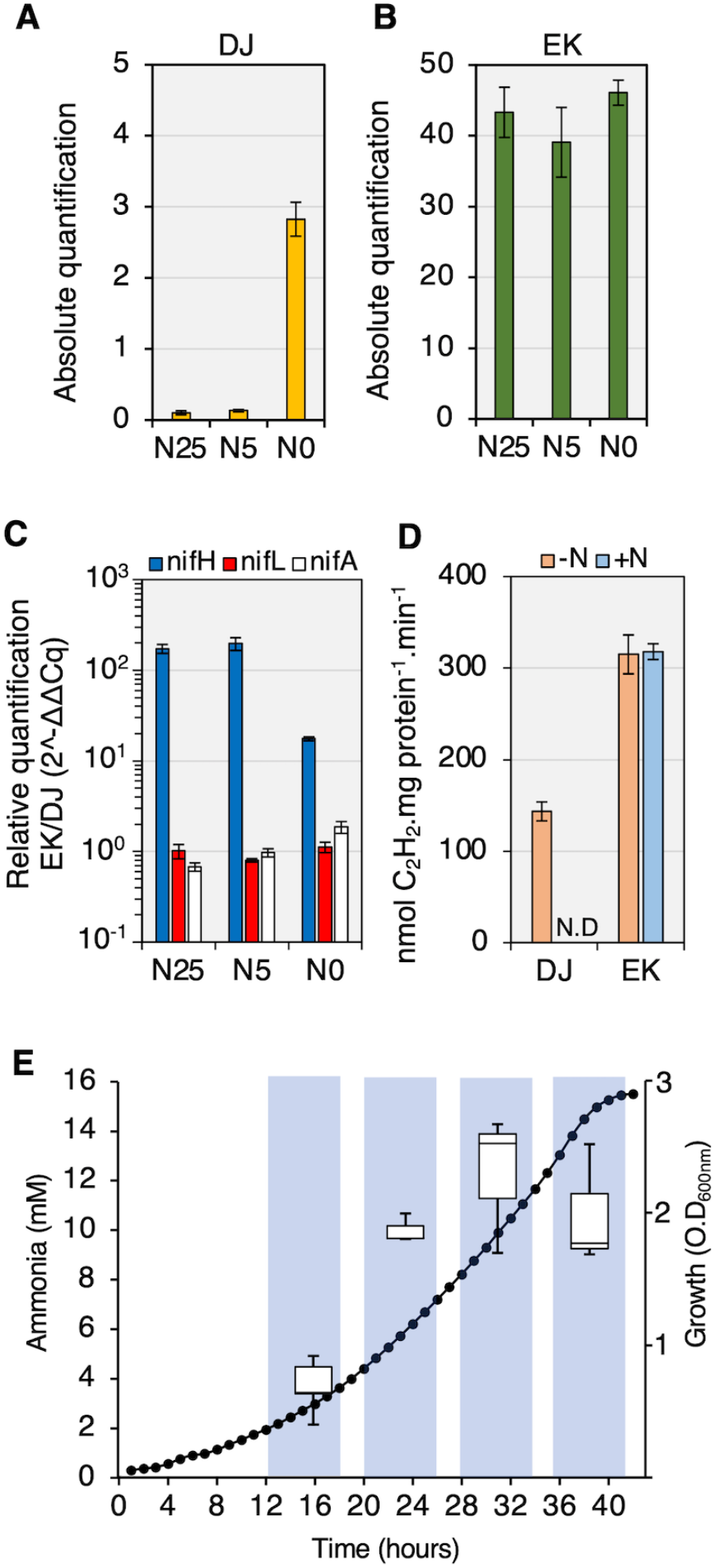
Constitutive expression and activity of nitrogenase results in ammonia excretion by the *nifA-E356K* strain (EK) with sucrose as the carbon source. (A) Absolute levels of *nifH* transcripts in the wild type (DJ) and (B) *nifA-E356K* (EK) under three different nitrogen regimes: 25 mM (N25), 5 mM (N5) or 0 mM (N0) ammonium acetate. (C) Relative levels of *nifH*, *nifL* and *nifA* transcripts between the strains EK and DJ. The graph is presented on a *log_10_* scale to emphasize that the relative levels of *nifL* and *nifA* transcripts are close to 1 in all conditions. (D) *In vivo* nitrogenase specific activities in the absence (-N) or presence (+N) of 20 mM ammonium chloride. Activity was determined by the acetylene reduction assay using cultures grown to an O.D_600nm_ between 0.3-0.4 as described in the methods. N.D: not detected. (E) Ammonia from the culture supernatant was quantified in the EK strain (left y axis, box blots) in the growth phases indicated by the bars shaded in blue (right y axis, closed circles).

When cultivated under diazotrophic conditions in nitrogen-free media supplemented with sucrose as carbon source, the EK strain excreted millimolar levels of ammonia (Fig. 1E). The onset of ammonia excretion occurred at the early stages of growth (O.D_600nm_ 0.3 - 0.6) reaching a peak at mid to late-exponential phase (O.D_600nm_ 1.4 - 2.0) and declined upon entry into stationary phase, potentially as a consequence of oxygen limitation and lower nitrogen fixation rates.

### Nitrogen fixation and ammonium assimilation are uncoupled in the *nifA-E356K* mutant strain

It is perhaps surprising that a single amino acid substitution in NifA enables ammonia excretion, since lowering the flux of ammonia assimilation through the glutamine synthetase-glutamate synthase (GS-GOGAT) pathway is expected to be an additional pre-requisite for high level release of ammonia. The enzyme glutamine synthetase (GS) is a key component for ammonia assimilation in proteobacteria and is subject to post-translational regulation via adenylylation in response to the nitrogen status (26, 27). Under excess nitrogen conditions, GS is adenylylated reducing its biosynthetic activity (3, 27). Comparison of the Mg^2+^, ATP-dependent glutamine synthetase biosynthetic (GSB) and the Mn^2+^, AsO4-, ADP-dependent glutamine transferase (GST) activities can provide a snapshot of GS adenylylation states *in vivo*, given that GSB activity can only be detected in the non-adenylylated enzyme subunits (28–30). We observed that GSB activity was higher in wild type (DJ) when compared to the mutant (EK) in all growth phases when cells were cultivated under diazotrophic conditions (green shaded plots in Fig. 2A). Conversely, GST activity was higher in the *nifA-E356K* strain (EK) than in the wild type (DJ) (Fig. 2B). A comparison of GST/GSB ratios in both strains (Fig. 2C) demonstrated that GS in the wild type (DJ) is likely to be entirely non-adenylylated under diazotrophic conditions (GST/GSB ratio ≦ 1), whilst the NifA-E356K mutant (EK) had much higher GST/GSB ratios (ranging from 15 to 30-fold) suggesting that in this mutant strain, GS is more heavily adenylylated. Under fixed nitrogen excess conditions (yellow shaded plots in Fig. 2), as anticipated, a marked reduction of GSB activity was observed in the DJ strain (Fig. 2A, yellow plot) followed by an increase in the GST activity (Fig. 2B, yellow plot). For the EK strain however, relatively minor changes in GSB and GST activities occurred in response to the nitrogen status. A slight increase in GSB activity was observed in the EK strain under nitrogen excess conditions, but this was not apparently associated with changes in the expression of GS (*glnA*) itself (Fig. S2). Taken together, these results confirm that the ability of the *nifA-E356K* mutant strain to assimilate ammonia via GS is reduced compared to the wild type under diazotrophic conditions. This is likely to be a consequence of increased adenylylation by the bifunctional adenylyl transferase enzyme GlnE, which carries out post-translational modification of GS in response to the nitrogen status. Since deletion of *glnE* prevents adenylylation of GS in *A. vinelandii* (31), we introduced a *glnE* deletion into the *nifA-E356K* background to generate the strain EKΔE. Under diazotrophic conditions, GSB and GST activities in the EKΔE strain were similar to the those in the wild type strain (DJ) and clearly distinct from the activities in the *nifA-E356K* strain (EK) (compare green shaded plots in Fig. 2A–B and Fig. 2D–E). However, as anticipated from the absence of adenylyl transferase activity, addition of ammonium to the media did not lead to either reduction of GSB or increased GST activity in the EKΔE strain when compared to the DJ strain (compare yellow shaded plots in Fig. 2A–B and Fig. 2D–E). As expected from the increased GS biosynthetic activity exhibited by the EKΔE strain under nitrogen excess conditions, ammonia excretion was ablated in this strain (Fig. 2F and Fig. S3). These results therefore imply that the ability of the *nifA-E356K* (EK) strain to excrete ammonia is dependent upon feedback regulation, resulting in constitutive adenylylation of GS and hence decreased ammonia assimilation.

**Fig. 2.**
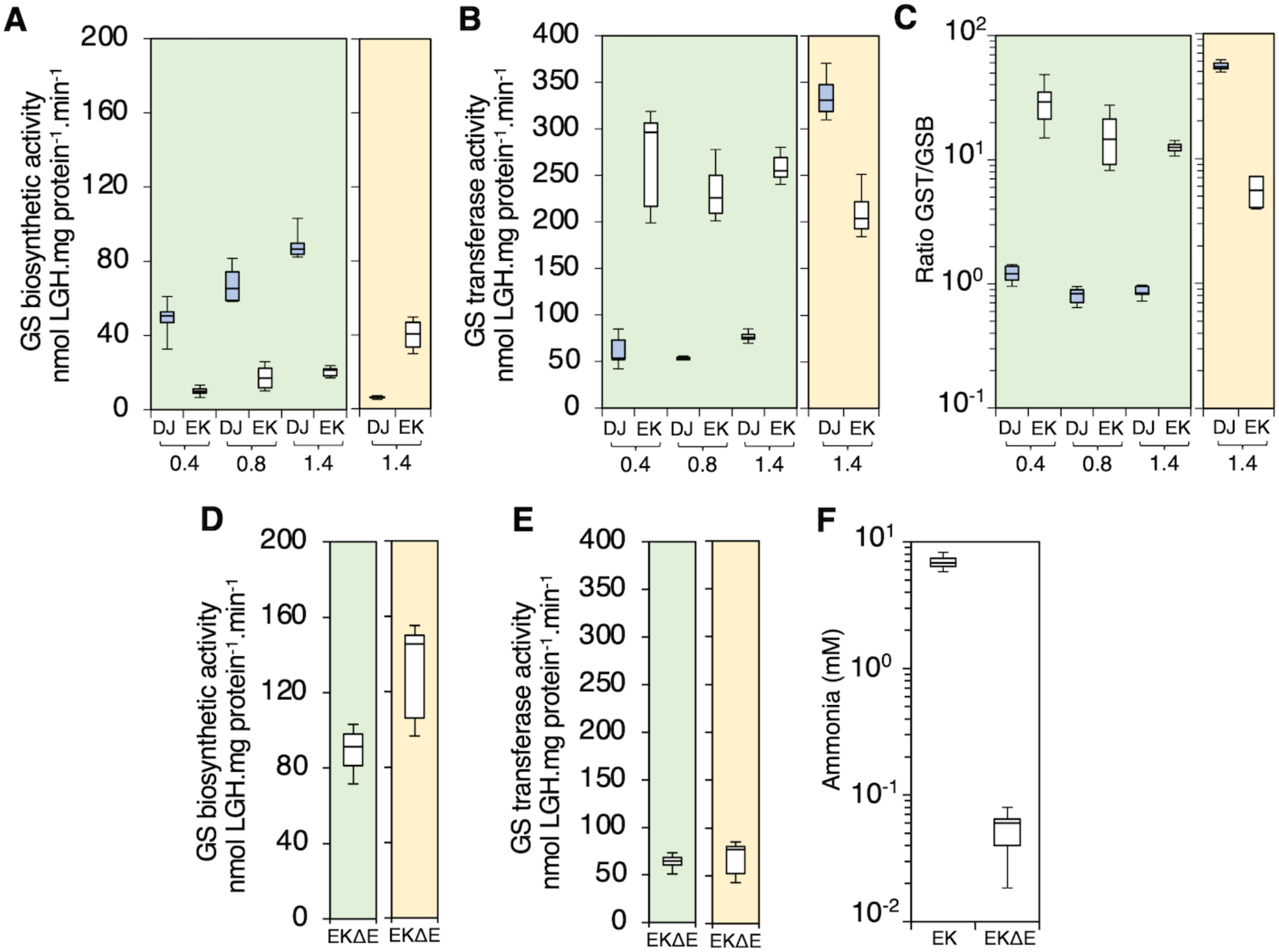
Ammonia excretion is dependent upon lower glutamine synthetase biosynthetic activity in the *nifA-E356K* strain (EK). (A) Glutamine synthetase (GS) biosynthetic and (B) transferase activities were measured in the wild type (DJ, blue box plots) and E356K (EK, white box plots) strains in three different phases of growth, corresponding to an O.D_600 nm_ of 0.4, 0.8 and 1.4 as indicated. Ratio between GS transferase (GST) and GS biosynthetic (GSB) activities is presented in (C) on a *log_10_* scale to emphasize that the ratio between GST and GSB activities are close to 1 in the wild type (DJ) in all growth phases. (D) Glutamine synthetase (GS) biosynthetic and (E) transferase activities measured in the EKΔE strain (*nifA-E356K* with a *glnE* deletion) at an O.D_600 nm_ of 0.8. The charts shaded in green represent the activities under diazotrophic conditions (-N), while those shaded in yellow represent the activity in the presence of excess fixed nitrogen, (20 mM NH_4_Cl, +N). (F) Ammonia from the culture supernatant was quantified in both EK and EKΔE strains grown under diazotrophic conditions.

### Carbon signalling dominates regulation of *nif* gene expression in the *nifA-E356K* strain

All the above experiments were conducted in media containing sucrose, a carbon source enabling a high flux through the TCA cycle that evidently maintains sufficient 2-oxoglutarate to activate NifA in *A. vinelandii*. Although the potential for carbon regulation, signalled via binding of 2-oxoglutarate to the NifA GAF domain, has been well established *in vitro,* the physiological relevance of 2-oxoglutarate in NifL-NifA regulation has not been clearly demonstrated *in vivo*. NifA-E356K requires 2-oxoglutarate in order to escape inhibition by NifL in the presence of GlnK *in vitro* (14, 21), suggesting that its ability to bypass nitrogen regulation *in vivo* might be regulated by carbon source availability. To facilitate correlation of nitrogenase activity with expression of the nitrogenase structural genes when strains were grown on different carbon sources we constructed strains containing a translational *nifH::lacZ* fusion located at a neutral site in the chromosome (see Supplementary Material and Table S1). Initial screening for the ability of the NifA-E356K protein to escape nitrogen regulation in several carbon sources (Figure S4) revealed that this variant protein supported strong activation of the *nifH* promoter under nitrogen excess conditions when grown in sucrose, glucose or glycerol. In contrast, only approximately 20-40% of the maximum activity was observed when cells were grown in succinate, fumarate, malate, pyruvate or acetate as the sole carbon source. Further comparisons of Av-NifA-E356K activity were performed comparing sucrose and acetate as carbon sources given that the growth penalty difference between the DJ and EK strains was significantly alleviated in acetate (Figure S5).

When wild type *A. vinelandii* was subjected to a carbon shift from sucrose to acetate, a significant reduction in both *nifH* expression (Fig. 3A) and nitrogenase activity (Fig. 3B) was observed and as expected, both activities were repressed in the presence of ammonium (+N). However, the ability of the EK strain to escape regulation by fixed nitrogen (+N) was severely compromised when grown on acetate (Fig. 3D and Fig. 3E). These results demonstrate that NifA-E356K responds to carbon status regulation *in vivo*, as anticipated from the *in vitro* characterization experiments (14, 21). Since 2-oxoglutarate is required to activate NifA only when NifL is present, we examined nitrogen and carbon regulation in the previously characterised strain AZBB163 in which *nifL* is disrupted by a kanamycin resistance cassette. (19). In this case, in contrast to NifA-E356K, nitrogenase activity was constitutive and not strongly influenced by the carbon source (Fig. S6B). However, this strain exhibited unexpected patterns of *nifH* expression when grown on sucrose that did not correlate with nitrogenase activity (Fig. S6A), potentially because *nifA* expression is not driven by the native *nifL* promoter in strain AZBB163 (19). Taken together, these results demonstrate that nitrogenase expression and activity is suppressed in the EK strain when grown under nitrogen excess conditions with acetate as the sole carbon source. Since the NifA-E356K protein is unable to escape nitrogen regulation mediated by NifL and GlnK when 2-oxoglutarate is limiting *in vitro*, this metabolite is likely to provide the physiological signal that triggers the carbon source response. Quantification of internal 2-oxoglutarate levels in strains grown on the different carbon sources (Fig. 3C and Fig. 3F) supports previous evidence that the levels of this metabolite are sensitive to the carbon and nitrogen supply (15, 32). In the wild type strain (DJ), 2-oxoglutarate levels dropped significantly in acetate compared to sucrose and a further decrease was observed when excess fixed nitrogen was present regardless of the type of carbon source (Fig. 3C). 2-oxoglutarate levels in the *nifA-E356K* strain were generally higher than in the wild type, but were influenced in a similar manner in relation to carbon and nitrogen source availability (Fig. 3F). Overall, the fluctuations in the 2-oxoglutarate levels correlated well with nitrogenase activity and expression for both the wild type (compare Fig. 3, panels A-C) and the *nifA-E356K* mutant (compare Fig. 3, panels D-F), reinforcing the importance of carbon signalling in the regulation of nitrogen fixation. Notably, when the 2-oxoglutarate level decreased below 350 μM in the *nifA-E356K* strain (Fig. 3F, acetate +N condition) nitrogen regulation was less effectively bypassed *in vivo* (Fig. 3, panels D-E), commensurate with previous *in vitro* biochemical data (14, 33). Consequently, lower levels of ammonia excretion were detectable when the *nifA-E356K* strain was grown on acetate (Fig. 3G). In accordance with this, the growth rate penalty observed in the EK (*nifA-E356K*) strain in the presence of sucrose was significantly reduced when acetate was the carbon source (Fig S5).

**Fig. 3.**
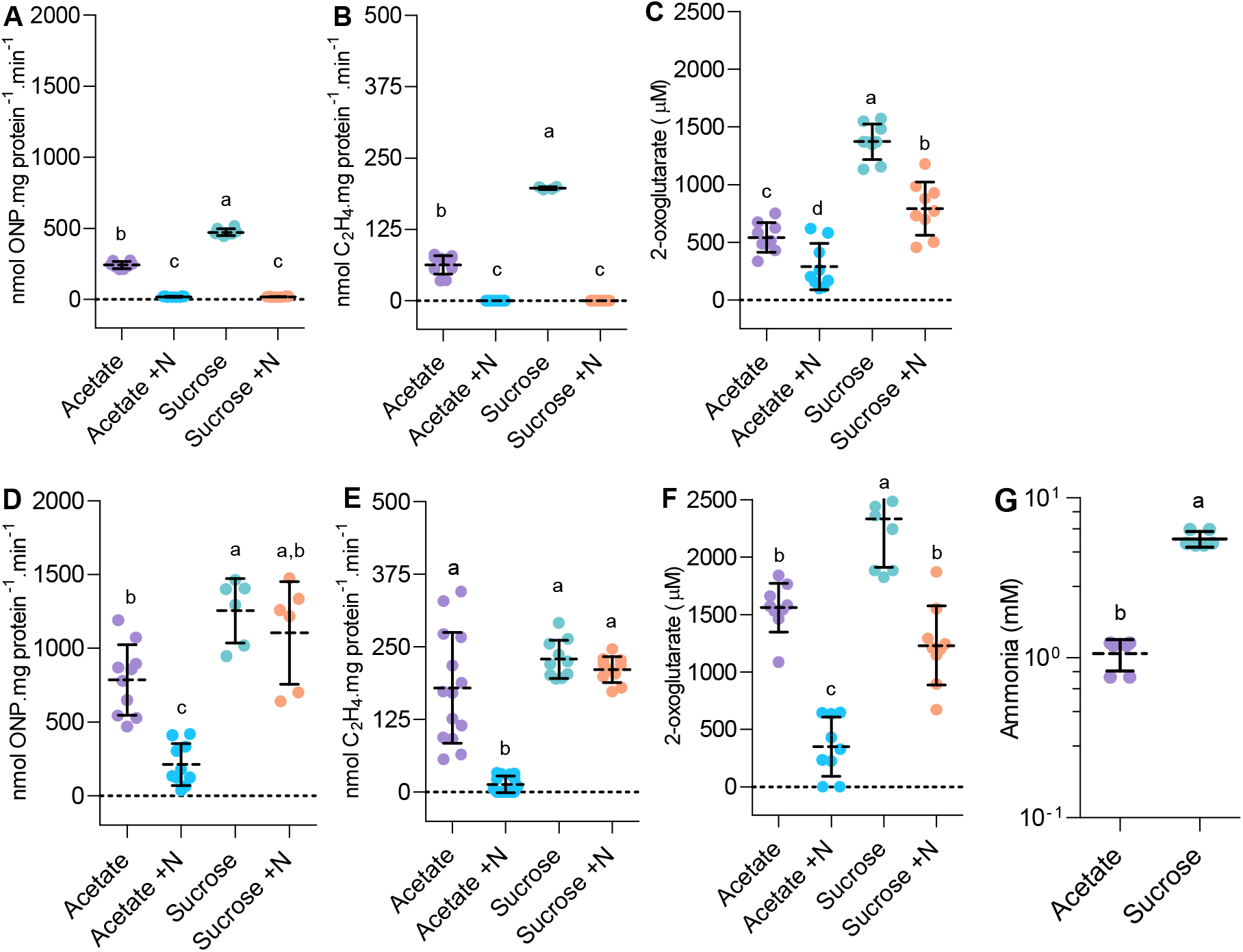
Carbon status regulation of nitrogenase expression and ammonia excretion in *A. vinelandii*. (A) Nitrogenase expression reported from a *nifH::lacZ* fusion, (B) nitrogenase activity and (C) internal 2-oxoglutarate levels are compared in the wild type (DJ) in acetate and sucrose in the absence (-N) or presence (+N) of 20 mM ammonium chloride. (D-F) The same comparisons as in (A-C) were done for the *nifA-E356K* strain (EK). For experiments in (A-F), strains were cultured to an O.D600nm between 0.2-0.3 in 30 mM acetate or 60 mM sucrose. To facilitate direct comparison of nitrogenase expression and activity, strain DJHZ was used in panels (A-B) whereas strain EKHZ was used in panels (D-E). These strains are isogenic to DJ and EK, respectively, except that they encode a *nifH::lacZ* fusion in the *algU* locus, a neutral site in the *A. vinelandii* genome (see Supplementary Material and Table S1). (G) Ammonia levels detected in the *nifA-E356K* (EK) strain when grown in either acetate or sucrose until cultures reached stationary phase. Plots followed by different letters are statistically different according to ANOVA with post-hoc Tukey’s HSD or a paired t-test in (G).

### NifA-E356K is a prototype for engineering conditional ammonia excretion in diazotrophic Proteobacteria

As the NifL-NifA operon is widely distributed in Proteobacteria (reviewed in 3) and the glutamate residue at position 356 in *A. vinelandii* NifA is highly conserved in other Proteobacteria (Fig. 4 panels A and B), we sought to evaluate if introduction of the reciprocal amino acid substitution in NifA proteins from other Proteobacteria, would yield the same regulation profile as in *A. vinelandii*. Using a previously established two-plasmid system to study the *A. vinelandii* NifL-NifA system in an *E. coli* background (11) we evaluated the activity of NifA variants from *Pseudomonas stutzeri* A1501 (22, 23) and *Azoarcus olearius* DQS-4 (34, 35). Both of these species are thought to be well adapted for the endophytic lifestyle and therefore are attractive model organisms for engineering ammonia excretion to benefit plant growth. The activities of wild type *P*. *stutzeri* NifA and *A*. *olearius* NifA in the presence of their corresponding NifL partners were higher than wild type *A. vinelandii* NifA under nitrogen-limiting conditions (-N) but as expected, were strongly inhibited in the presence of fixed nitrogen (+N) (Fig. 4, panels C-E). In contrast, the reciprocal NifA-E356K substitutions in *P. stutzeri* NifA (Ps-NifA-E356K) and *A. olearius* NifA (Ao-NifA-E351K) gave rise to constitutive activation of the *nifH* promoter in nitrogen replete (+N) conditions in *E. coli* (Fig. 4 panels D-E, respectively).

**Fig. 4.**
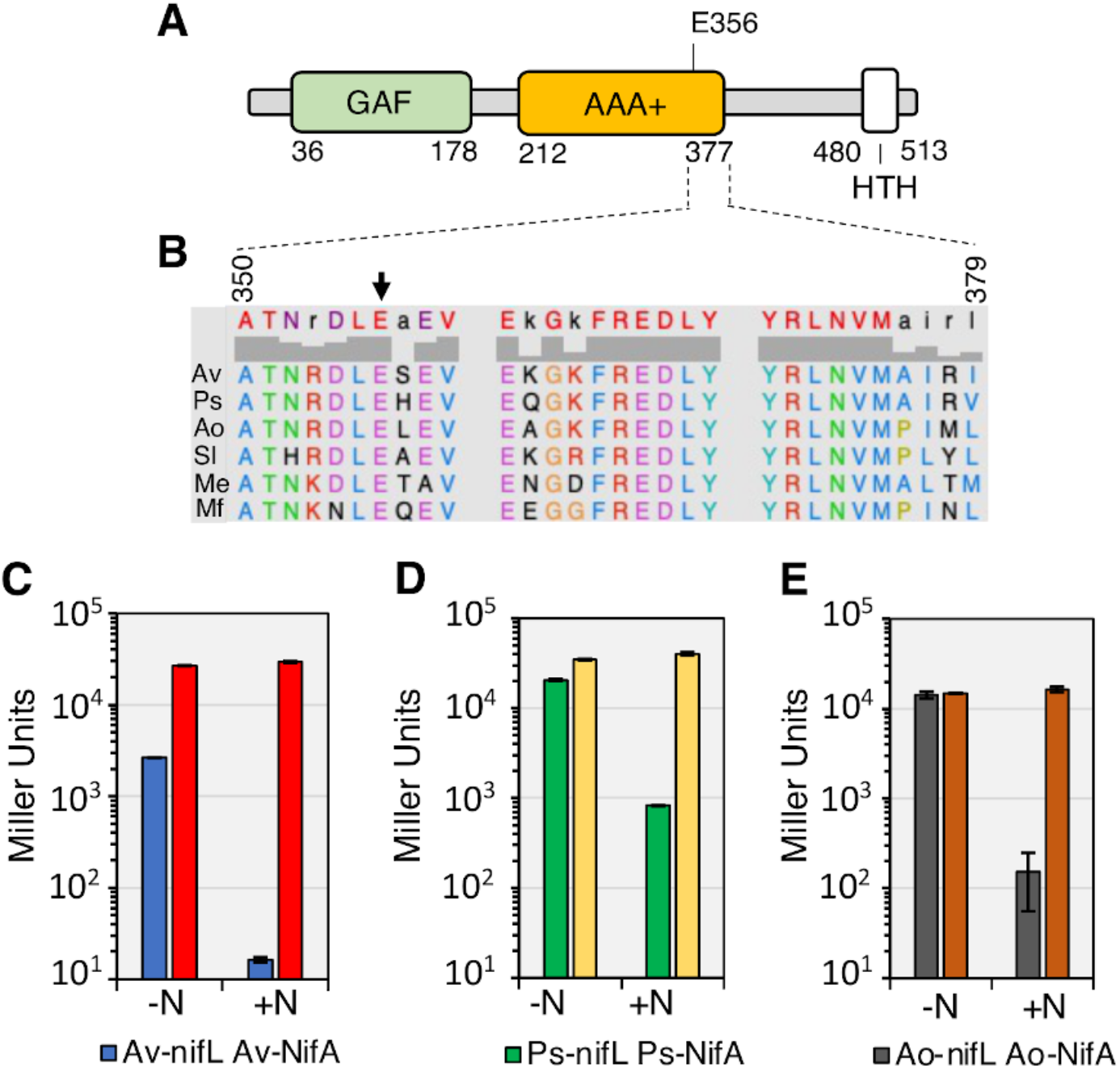
Reciprocal amino acid changes (related to *nifAE356K*in *A. vinelandii*) yield constitutively active NifA in Proteobacteria. (A) Diagram of the *A. vinelandii* NifA domains. (B) Alignment of residues close to E356 (black arrow) in the central AAA+ domain of NifA proteins. Sequence numbers refer to *A. vinelandii* NifA. Sequences used in the alignment are Av: *A. vinelandii* DJ, Ps: *Pseudomonas stutzeri* A1501 (Gammaproteobacteria), Ao: *Azoarcus olearius* DQS4, Sl: *Sideroxydans lithotrophicus* ES-1 (Betaproteobacteria), Me: *Martelella endophytica* YC6887 (Alphaproteobacteria) and Mf: *Mariprofundus ferrooxydans* M34 (Zetaproteobacteria). Panels C to E show *β*-galactosidase activities in the *E. coli* ET8000 chassis resulting from activation of a *nifH::lacZ* fusion (plasmid pRT22) by wild type and variant NifL-NifA systems from three different diazotrophs. Plasmids used to express NifL-NifA variants are as follows. (C) pPR34: Av-NifL-NifA, pPMA: Av-NifL-NifA-E356K; (D) pMB1804: Ps-NifL-NifA, pMB1805: Ps-NifL-NifA-E356K; (E) pMB1806: Ao-NifL-NifA, pMB1807: Ao-NifL-NifA-E351K. The assays were performed in NFDM media supplemented with 2% glucose in either nitrogen-limiting (200 μg/ml of casein hydrolysate, -N) or nitrogen excess (7.56 mM ammonium sulphate, +N) conditions.

To examine the properties of the Ps-NifA-E356K substitution in its endophytic host, we introduced the corresponding *nifA* mutation into the chromosome of *P. stutzeri* A1501. However, contrary to *A. vinelandii,* this single mutation (in strain Ps-EK) did not result in constitutive *nifH* transcription (Fig. S7), presumably because expression of the *nifLA* operon itself is regulated by nitrogen availability in *P. stutzeri* (23, 36). In order to remove this second layer of nitrogen regulation, we replaced the native *P.stutzeri nifL* promoter with the *A. vinelandii nifL* promoter. Although this replacement (in the strain Ps_nifLA^C^) suppressed nitrogen control of *nifA* transcription as anticipated, constitutive *nifH* transcription was not observed, confirming that the *P. stutzeri* wild-type NifL-NifA system remains responsive to nitrogen regulation when expressed constitutively (Fig. S7). To examine the intrinsic ability of Ps-NifA-E356K to escape nitrogen control, we combined the *A. vinelandii nifL* promoter replacement with the *nifA-E356K* mutation in *P. stutzeri* (Fig. 5A). Although this strain (Ps-EK^C^) expressed relatively low levels of *nifLA* transcripts under diazotrophic (-N) conditions (Fig. S7 F-G), direct correlation between the levels of *nifLA* and *nifH* transcripts was observed in excess nitrogen (+N) conditions, confirming that the E356K substitution enables Ps-NifA to escape nitrogen regulation mediated by NifL and GlnK in *P. stutzeri* (Fig. S7, panels F-H).

**Fig. 5.**
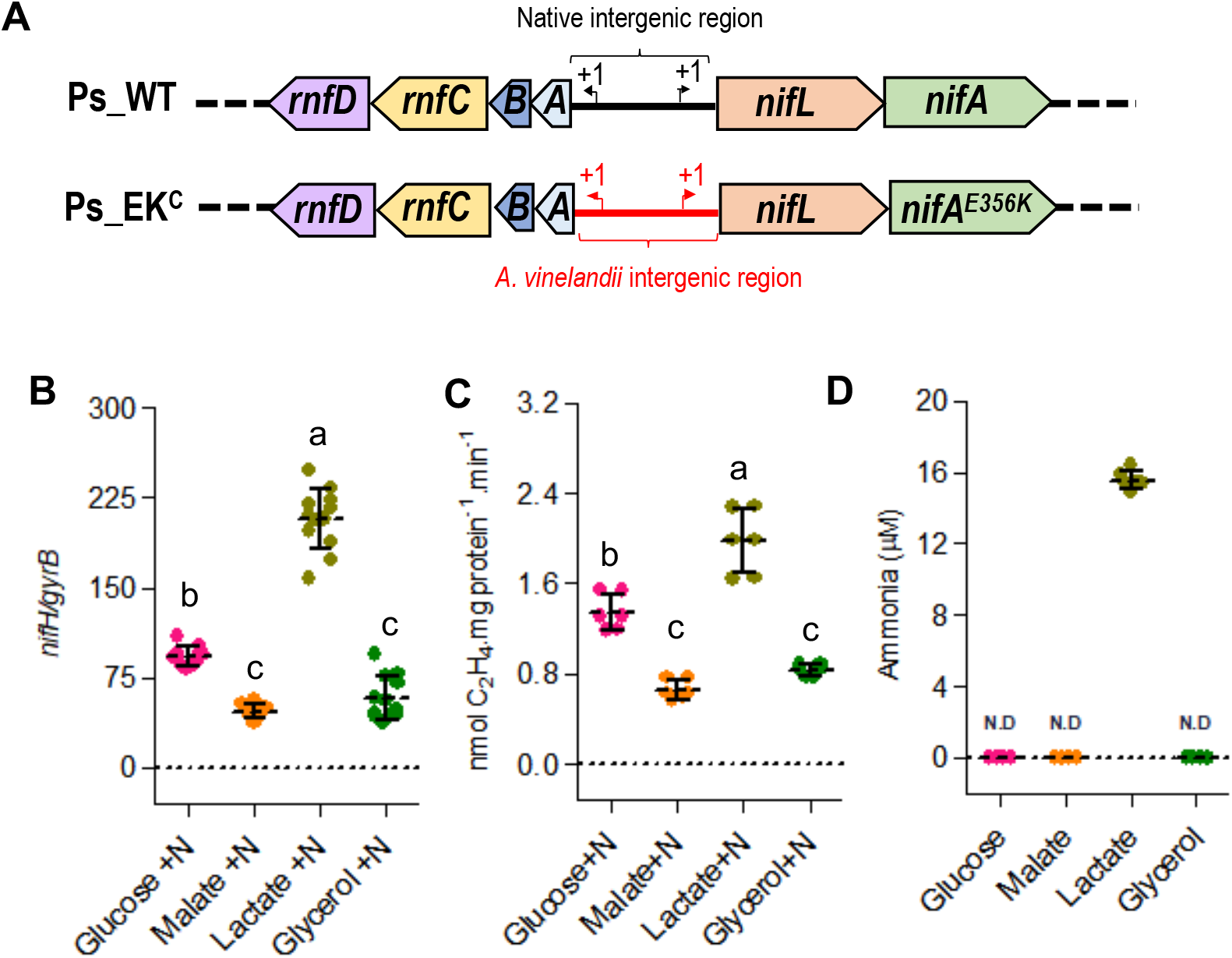
The Ps-NifA-E356K protein is able to escape NifL inhibition in *P. stutzeri* leading to carbon-dependent ammonia excretion. (A) Diagram depicting the modifications in the *P. stutzeri nifA-E356K* strain (Ps_EKc) compared to wild type *P. stutzeri* (Ps_WT). Drawings are not to scale. (B) Levels of *nifH* transcripts in the *P. stutzeri* Ps_EK^c^ strain in the presence of 5 mM NH_4_Cl (+N). (C) Nitrogenase activity in the *P. stutzeri* strain (Ps_EK^c^) in the presence of 5 mM NH_4_Cl (+N). (D) Comparison of ammonia excretion profiles in different carbon sources. In all cases, the Ps_EKc strain was grown in UMS-PS supplemented with 30 mM glucose, 45 mM malate, 60 mM lactate or 60 mM glycerol (to provide balanced carbon equivalents). N.D: not detected. Plots followed by different letters are statistically different according to ANOVA with post-hoc Tukey’s HSD.

The Ps-EK^C^ strain was able to activate *nifH* transcription on a variety of carbon sources when grown under nitrogen excess conditions (+N) with lactate yielding the highest level of *nifH* activation followed by glucose, malate and glycerol (Fig 5B). This carbon source-dependent activation of *nifH* transcription by Ps-NifA-E356K correlated directly with the level of nitrogenase activity in each condition (compare Fig. 5B and 5C). As anticipated from the relatively high level of *nifH* transcription and nitrogenase activity conferred by growth on lactate, ammonia excretion was only observed when the Ps-EK^C^ strain was grown on this carbon source (Fig. 5D). Finally, we observed no growth penalty for the Ps-EK^C^ strain when grown on complex media (LB) or in minimal media supplemented with glucose, malate, lactate or glycerol as carbon sources (Fig S8), which implies that the relatively moderate activation of *nif* gene transcription in the Ps-EK^C^ strain, allows carbon regulated ammonia excretion without severe impacts to bacterial fitness. Altogether, these observations suggest that introducing the reciprocal E356K substitution into NifA proteins from diazotrophic Proteobacteria, may be broadly applicable for engineering new bacterial strains with carbon-controlled excretion of ammonia.

## Discussion

In order to cope with the energetic cost of biological nitrogen fixation, diazotrophic bacteria require sophisticated signal transduction mechanisms ensuring efficient adaption to changing conditions whilst successfully competing in the environment. Achieving an appropriate balance between carbon and nitrogen metabolism is particularly challenging for organisms that fix nitrogen, requiring diversion of ATP and reducing equivalents from central metabolism to ensure nitrogenase catalytic rates that meet the nitrogen demands required for growth. Hence the ability to sense carbon availability in addition to the nitrogen status, is paramount to resource allocation and to resolve conflicting metabolic demands.

The physiological signal for carbon status control is most likely to be 2-oxoglutarate given the correlation observed here between the level of this metabolite with nitrogen regulation *in vivo*, together with our previous biochemical demonstration of the importance of this ligand in NifL-NifA regulation (14, 37). We propose that this additional level of metabolite regulation provides a mechanism to integrate signals of the carbon and nitrogen status to ensure that sufficient carbon resources are available to support diazotrophy. The *in vitro* data indicate that when 2-oxoglutarate is limiting, NifL, forms a binary complex with NifA, which inhibits its activity, even under nitrogen-limiting conditions when GlnK is fully uridylylated and unable to interact with nifL (Fig. 6A). However, when sufficient levels of 2-oxoglutarate are available (Fig. 6B) the NifL-NifA complex dissociates, enabling NifA to activate *nif* transcription (2, 4, 14, 37). Upon a switch to excess nitrogen conditions, GlnK becomes de-uridylylated allowing the formation of a ternary GlnK-NifL-NifA complex that inactivates NifA irrespective of the level of 2-oxoglutarate (Fig. 6C and 6D). Hence in the wild-type NifL-NifA system, the nitrogen status signal overrides the metabolic signal of the carbon status, when excess fixed nitrogen is available. In contrast, in the variant NifA-E356K protein studied here, the integration between nitrogen and carbon control is disrupted. Although this substitution in the AAA+ domain of NifA (red star, Fig. 6E) perturbs the interaction with NifL, the GlnK-NifL-NifA-E356K ternary complex still forms if 2-oxoglutarate is limiting (Fig. 6E). In contrast, when 2-oxoglutarate levels are sufficient, conformational changes triggered by its binding to the GAF domain disrupt the ternary complex, enabling NifA-E356K to be active in the presence of excess fixed nitrogen (Fig. 6F). Therefore, although the E356K substitution escapes nitrogen control, conferred by resistance to the GlnK bound form of NifL, this is contingent upon the binding of 2-oxoglutarate to the GAF domain of this variant protein. The response of the NifL-NifA system to 2-oxoglutarate thus emphasises the key role of this metabolite as a master signalling molecule (15). Consequently, the ability of the NifA-E356K variant to bypass nitrogen regulation *in vivo* in both *A. vinelandii* and *P. stutzeri* is dependent on the carbon status. The crucial role of carbon-mediated signalling in the regulation of nitrogen fixation was evident from reduced *nifH* transcripts and activity of nitrogenase when the *A. vinelandii nifA-E356K* strain was cultured under nitrogen excess conditions with acetate as sole carbon source, which correlated with a significant decrease in the level of 2-oxoglutarate and a 6-fold reduction in ammonium excretion compared with sucrose as carbon source. Similarly, in *P. stutzeri* where lactate appears to be a preferred carbon source to support nitrogen fixation in comparison to glucose, the *nifA-E356K* mutant exhibited the highest level of *nifH* transcripts and nitrogenase activity under nitrogen excess conditions when lactate was provided as a carbon source. Not surprisingly, amongst the carbon sources tested, the *P. stutzeri* EK^C^ strain only excreted ammonia when provided with lactate under our experimental conditions. The capacity for runaway expression of *nif* genes, constitutive nitrogenase activity and ammonia excretion is therefore dependent on the nature of the carbon source. Our studies with *A. vinelandii* and *P. stutzeri* therefore demonstrate the potential to exploit the intrinsic carbon-sensing mechanism of the NifL-NifA system to provide conditional release of fixed nitrogen and hence alleviate the fitness penalty associated with constitutive expression of nitrogenase.

**Fig. 6.**
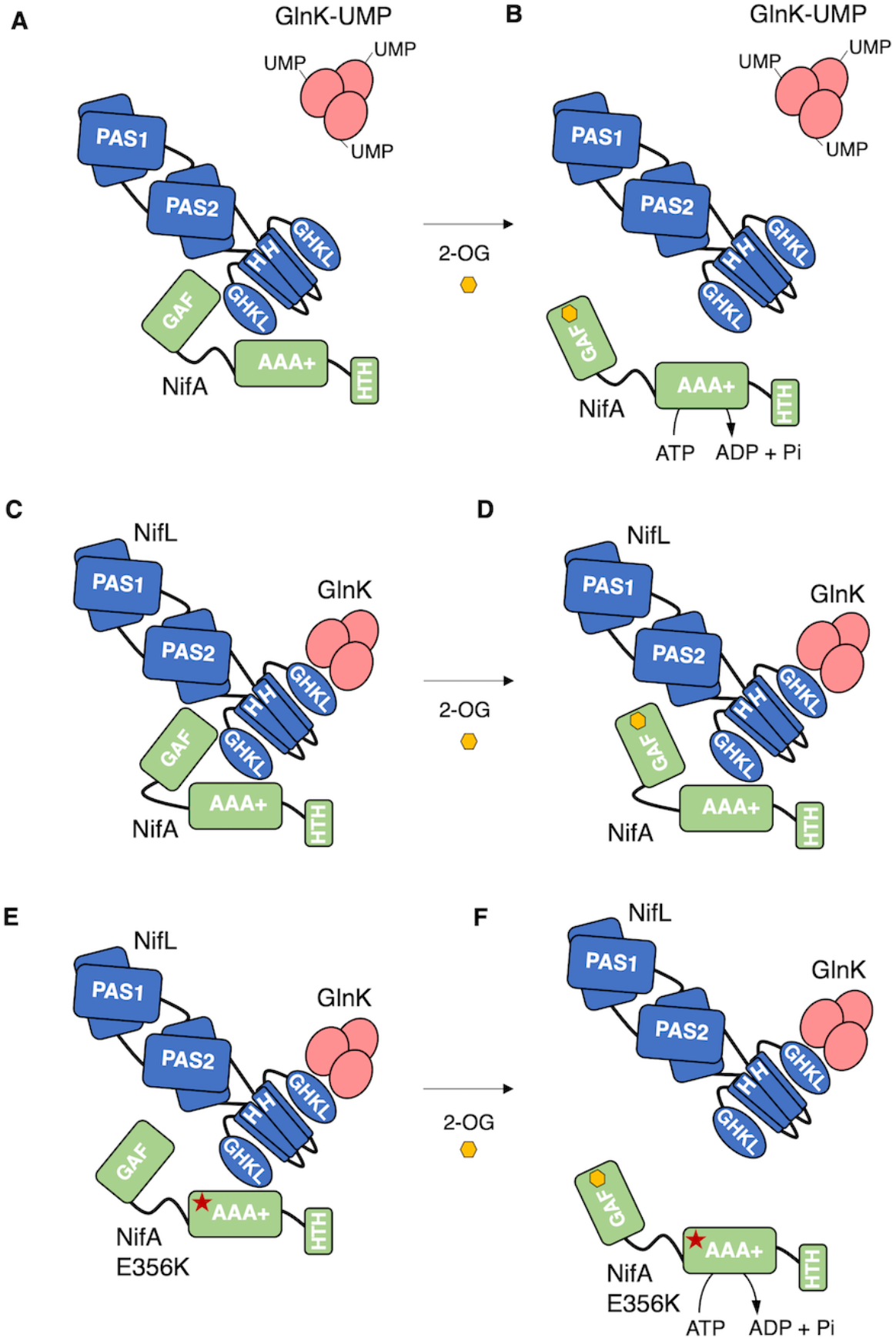
Model for 2-oxoglutarate regulation of NifA activity based on genetic and biochemical experiments. (A) When 2-oxoglutarate levels are low, NifL can inhibit NifA even under nitrogen limiting conditions when GlnK is uridylylated (GlnK-UMP) and unable to interact with NifL. (B) Binding of 2-oxoglutarate (2-OG, yellow hexagon) to the GAF domain of NifA, disrupts the binary NifL-NifA interaction activating NifA. s(C) Under nitrogen excess conditions, non-covalently modified GlnK, interacts with the GHKL domain of NifL, stimulating the formation of a ternary complex between GlnK, NifL and NifA that inhibits NifA activity. (D) The GlnK-NifL-NifA ternary complex is stable even if the GAF domain in NifA is saturated with 2-OG. (E) The E356K substitution in the AAA+ domain of NifA (red star), perturbs the interaction with NifL in the absence of the non-modified form of GlnK. However, under nitrogen excess conditions, the GlnK-NifL-NifA-E356K ternary complex is formed when 2-OG in limiting. (F) When 2-OG levels are sufficient, conformational changes triggered by 2-OG binding to the GAF domain disrupt the ternary complex, activating NifA-E356K. Physiologically, interactions depicted in (E), may arise upon growth *in vivo* under carbon limiting conditions. A switch to a preferred carbon source, will lead to an increase in 2-OG, triggering the conformation change that allows NifA-E356K to escape from GlnK-NifL inhibition (F).

Bypassing nitrogen regulation of the NifL-NifA system to activate constitutive expression of nitrogenase would not by itself be anticipated to promote ammonia release, if the excess ammonia can be assimilated by the GS-GOGAT pathway. We have demonstrated that overexpression of nitrogenase in the *A. vinelandii nifA-E356K* strain leads to feedback regulation of GS activity via co-valent modification by the adenylyl transferase activity of GlnE. This reduction of nitrogen assimilation via post-translational modification of glutamine synthetase is a key factor in enabling ammonia excretion, which is not observed in the *nifA-E356K* strain when the *glnE* gene is deleted. Hence, in *A. vinelandii*, ammonia excretion is also dependent on the native feedback regulation of GS activity, exacerbated by higher rates of nitrogen fixation in the *nifA-E356K* strain when grown on a carbon source that sustains high levels of 2-oxoglutarate under excess nitrogen conditions. This uncoupling of nitrogen fixation from ammonium assimilation is somewhat analogous to what is observed in differentiated nitrogen-fixing bacteroids in the legume-rhizobium symbiosis, where the flux through the ammonia assimilation pathway is severely restricted to enable release of most of the nitrogen fixed by the symbiont (38, 39). Analogous strategies to decrease the activity of GS in non-symbiotic bacteria have resulted in ammonia excretion (40–42), but to date have not been combined with mutations that express high levels of nitrogenase on a conditional basis as deployed here.

Since introduction of the reciprocal E356K substitution into NifA proteins from other members of the Proteobacteria also results in nitrogen-insensitive activators when analysed either in a heterologous chassis, *E. coli* (Fig. 4), or in *P. stutzeri* (Fig. 5), this strategy may allow generation of new diazotrophic strains with conditional excretion of ammonia. The carbon responsive control mechanism could present an opportunity for activation of ammonium excretion contingent upon carbon sources provided by root exudates of crops, a feature highly desirable in the engineering of a synthetic symbiosis (43, 44). However, given the regulatory complexities associated with fine-tuning nitrogen regulation in diverse Proteobacteria, additional manipulations to disrupt transcriptional control of NifL-NifA expression itself or the coupling between nitrogen fixation and assimilation may be needed to achieve ammonia excretion (3, 45). One such example explored in this study was the need to remove native nitrogen regulation from the *P. stutzeri* A1501 *nifLA* promoter. Serendipitously, we demonstrated that providing relatively low levels of the *nifA-E356K* transcripts in *P. stutzeri* generated a strain with carbon-regulated ammonia excretion without the severe growth penalties observed in *A. vinelandii*. Furthermore, we demonstrated that the levels of ammonia excretion are directly correlated with specific nitrogenase activity rates in both organisms analysed. Under optimal conditions of carbon and oxygen supply, the *A. vinelandii nifA-E356K* mutant sustained a very high rate of nitrogenase activity (200-300 nmol C_2_H_4_.mg protein^-1^.min^-1^) under nitrogen excess conditions. On the other hand, in *P. stutzeri* the levels of nitrogenase activity conferred by *nifA-E356K* were at least 100-fold lower under our experimental conditions. Hence, *A. vinelandii* excreted millimolar levels of ammonia in contrast to the micromolar levels observed in *P. stutzeri*. As the ammonium excreting strain from *P. stutzeri* was not subject to the same growth penalty observed in the *A. vinelandii* counterpart, we anticipate that these studies will guide future efforts to define more precise trade-offs to engineer nitrogen releasing strains that do not have a competitive disadvantage in the rhizosphere. Moreover, the addition of multi-layered regulatory control of ammonia excretion by expressing activator variants under the control of promoters that respond to specific signalling molecules exchanged between the plant and the bacteria, may deliver the required level of specificity for the establishment of an efficient synthetic symbiosis (46, 47).

## Materials and Methods

Detailed methods are available in the Supplementary Material. Bacterial stains are listed in Table S1, plasmids are listed in Table S2 and primers are listed in Table S3.

## Author Contributions

M.B.B, Y-P.W and R.D designed research, M.B.B, P.B. and C.A-A. performed experiments and M.B.B, Y-P.W and R.D. wrote the manuscript

## Acknowledgements

We are indebted to Brett Barney for the strain AZBB163 and to Adriano Stefanello for the plasmid pAAS1544. We are grateful to all members of the laboratory support team at the John Innes Centre for their excellent assistance. This study was supported by the UKRI Biotechnology and Biological Sciences Research Council (grants BB/N013476/1 and BB/N003608/1), the Royal Society (ICA\R1\180088), the National Science Foundation of China (NSFC, grant 32020103002), and the National Key R&D Program of China (grant 2019YFA0904700).

