## Supplementary Material for "Disrupting hierarchical control of nitrogen fixation enables carbon-dependent regulation of ammonia excretion in soil diazotrophs"

This file includes:

Materials and Methods

Supplementary Figures S1-S8

Supplementary Tables S1-S3

Supplementary references

**Materials and Methods**

**Bacterial strains and growth conditions.** The bacterial strains used in this study are listed in Table S1. For routine procedures *E. coli* strains were grown at 37°C in LB medium (1) or in NFDM (2) for β-galactosidase activity assays. Media for *E. coli* ST18 (3) was supplemented with 50 µg/mL of ALA (5-aminolevulinic acid) unless counter-selection was required. *A. vinelandii* was grown at 30°C and 250 rpm in NIL medium (containing 0.2 g/L $MgCl_2$, 90 mg/L $CaCl_2$, 0.8 g/L $KH_2PO_4$, 0.2 g/L $K_2HPO_4$, 14 mg/L $Na_2SO_4$, 120 mg/L $Fe_2(SO_4)_3$ and 2.4 mg/L $Na_2MoO_4$) supplemented with either 2% sucrose (approximately 60 mM) as described previously (4, 5) unless stated otherwise. Alternatively, as the NIL medium was prone to precipitation hampering efforts to automate the measurement of growth rate parameters, a novel minimal medium, hereafter named MBB (*M*inimal *B*acterial *B*roth), was used for automated growth experiments on a plate reader. MBB contained 0.6 g/L $K_2HPO4$, 0.4 g/L $KH_2PO4$, 1.1 g/L NaCl, 0.4 g/L $MgSO_4$, 20 mg/L $CaCl_2.2H_2O$, 10 mg/L $MnSO_4.H_2O$, 65.6 mg/L Fe(III)-EDTA, 2 mg/L $Na_2MoO_4.2H_2O$ and 2% (20 g/L) sucrose. No apparent differences in the growth rate, ammonium excretion or diazotrophy (ability to grow on atmospheric $N_2$), were observed comparing NIL and MBB media when 2% sucrose was used as carbon source. Therefore, implementation of MBB allowed automation of growth parameter measurements, without detrimental effects to physiological traits of interest. *P. stutzeri* was grown at 30°C in LB for routine pre-inocula and for conjugations. For experiments requiring defined media such as measurement of nitrogenase activity, *P. stutzeri* was grown in a modified version of the UMS medium (6), hereafter named UMS-PS. UMS-PS was obtained by supplementing the UMS medium with 1:1000 dilution of vitamin solution (1 g/L thiamine hydrochloride, 2 g/L D-pantothenic acid, 1 g/L nicotinic acid, and 0.1 g/L biotin) and 50 mL/L of Kalininskaya phosphate buffer (KP buffer – $K_2HPO_4$ 33.4 g/L, $KH_2PO_4$ 17.4 g/L (7)), which significantly helped to reduce flocculation. The carbon source used for *P. stutzeri* was 2% glucose unless stated otherwise. 5-10 mM of $NH_4Cl$ was added to UMS-PS for non-diazotrophic growth as indicated. Antibiotics were used as follows: carbenicillin

50 µg/mL (*E. coli*), chloramphenicol 15 µg/mL (*E. coli*), tetracycline 5 µg/mL (*E. coli A. vinelandii* and *P. stutzeri*), kanamycin 50 µg/mL (*E. coli* and *P. stutzeri*) and 1-3 µg/mL (*A. vinelandii*), trimethoprim 15 µg/mL (*E. coli*) and 90 µg/mL (*A. vinelandii*).

**Preparation of *A. vinelandii* competent cells and transformation.** Competent cells of *A. vinelandii* were obtained in molybdate and iron depleted competence medium essentially as in (5), except that the medium was amended with 7 mM of $MgCl_2$. In brief, strains streaked for single colonies in competence medium plates were incubated at 30°C until a green fluorescent siderophore was produced (5-7 days), indicative of competence (5, 8). Subsequently, a loopful of cells was resuspended in 500 µL of P-buffer (4.6 mM $K_2HPO_4$,1.5 mM $KH_2PO_4$) and then mixed with 250-500 ng of a linear DNA fragment carrying the desired mutation. This mixture was then spotted onto the centre of a competence medium plate and further incubated overnight at 30°C. On the following day, cells were scraped off the plates and resuspended into 1 mL of P-buffer as above. A serial dilution was then spread onto NIL medium supplemented with 25 mM ammonium acetate with antibiotics to obtain single colonies. In the case of the *nifA-E356K* mutant, recombinant colony selection was achieved by recovery of diazotrophy (ability to grow on atmospheric $N_2$) using competent cells from a *nifA* deletion strain (DJA), without antibiotic selection, to generate the strain EK. Given that *A. vinelandii* can accumulate up to 80 copies of its chromosome under certain conditions (5, 9) newly recombinant colonies were exhaustively streaked on selective media (20 times or more) to ensure efficient chromosome segregation and homogeneity of the mutant genotype.

**Construction of *A. vinelandii* mutants.** To construct the *A. vinelandii nifA-E356K* mutant strain (EK), a fragment of 1569 pb corresponding to the *A. vinelandii nifA* gene carrying discrete base pair changes, GAA->AAG (yielding the E356K substitution), was amplified by PCR from the plasmid pAAS1544 (Table S2) with primers ASS-3 and ASS-55 (Table S3). The above base change also introduced a recognition site for the restriction enzyme, AcuI, which assisted genotypic screening. This PCR fragment was then directly transformed into a *nifA* deletion background and recombinant colonies were selected on the basis of diazotrophy recovery as described above. To construct mutants in the structural nitrogenase gene (*nifH*), a fragment of 1874 pb corresponding to the *nifHD* region (position 136301 - 138174) was amplified with primers 2-45 and 2-46 and cloned as a BamHI/HindIII fragment into pBlueScript II KS +, yielding pMB1724. Subsequently, the *tetA* gene (tetracycline resistance) was PCR amplified from pALMAR3 (primers 2-43 and 2-44) and cloned into the BglII and EcoRI sites of pMB1724 to yield pMB1725. This plasmid (Δ*nifH::tetA*), was linearized with ScaI and transformed, as described above, into *A. vinelandii* wild type (DJ) and *nifA-E356K* (EK) to generate the strains DJH and EKH, respectively. Construction of the *glnE* gene deletion strain (EKΔE), which carries a trimethoprim (*tmp*) resistance marker was performed by transformation of a linear PCR product obtained with primers M13F(-47) and M13R(-48) from the plasmid pMB1840. This plasmid, was obtained by isothermal assembly of fragments upstream (906 bp, position 4547284 - 4548189, primers 7-28/7-29) and downstream (752 bp, position 4543778 - 45444529, primers 7-32/7-33) of *glnE*, with a 631 bp fragment, obtained from pUC18T-mini-Tn7T-Tp with primers 7-30/7-31, encoding the *tmp* resistance gene, into pk18mobsacBKm linearized with SmaI. *A. vinelandii* reporter strains were obtained by insertion of a *nifH::lacZ* fusion into the *algU* locus. In *A. vinelandii* (DJ), the *algU* gene is naturally inactivated by an insertion sequence yielding the non-gummy phenotype (10, 11). We therefore anticipated that this was a convenient locus to insert a *nifH::lacZ* reporter whilst keeping the original *nifHDK* locus intact and avoiding problems that may arise from plasmid instability. This strategy facilitated comparison of both nitrogenase and NifA activities in a single *A.vinelandii* strain background. To obtain the *nifH::lacZ* reporter strains, the plasmid pMB1816 was linearized with ScaI and transformed into *A. vinelandii* as described above. pMB1816 was constructed by isothermal assembly of a 409 bp fragment corresponding to the *nifH* promoter (position 136401 – 136809, primers 5-49/5-50) with a 3057 bp (primers 5-46/5-464) and the *lacZ* gene derived from pRT22 into an *algU* integrative plasmid backbone. The latter was obtained by assembly of fragments upstream (795 bp, position 1329654 – 1330448) and downstream (813 bp, position 1332074 – 1332886) of the *algU* gene with an 1811 bp fragment corresponding to the *ori* and ampicillin resistance gene from pUC19 and a 631 bp fragment carrying the trimethoprim resistance gene.

**Construction of *P. stutzeri* mutants.** Mutagenic suicide plasmids based on the pk18mobsacBKm vector were conjugated into *P. stutzeri* essentially as described (12) by mixing recipient and donor strains in two proportions (50:1 and 10:1), except that the *E. coli* strain ST18 (3) was used as donor and that the whole procedure was performed on LB-agar medium. The *E. coli* ST18 was counter-selected after bi-parental mating based on its auxotrophy to ALA (5-aminolevulinic acid). Selection of double crossovers was performed in LB-agar supplemented with 10% sucrose. The *P. stutzeri* strain Ps_nifLA$^{C}$, was obtained by replacing the native *rnf-nifLA* intergenic region by the reciprocal region from *A.vinelandii* using pMB2005. This plasmid was obtained by fusing the *rnf-nifLA* intergenic region (444 bp; primers 8-19/8-20) from *A. vinelandii* downstream to a fragment of the *P. stutzeri rnfAB* genes (1160 bp; primers 8-17/8-18) and upstream to a fragment of the *P. stutzeri nifL* gene (1638 bp; primers 8-21/8-22) using isothermal assembly into pk18mobsacBKm linearized with SmaI. To generate the *P. stutzeri* strain Ps_EK$^{C}$ (See Fig. 5A) the plasmid pMB2006 was used to the generate the strain Ps_EK$^{C}$-*tetA*. Subsequently the *tetA* resistance cassette was recovered from the genome using pMB2007. The plasmid pMB2006 was obtained by fusing 847 bp downstream Ps-*nifA* (primers 8-2/8-1) with a 2364 bp fragment from the Ps-*nifLAE356K* (primers 8-3/8-11b; from pMB1805) by isothermal assembly into pk18mobsacBKm linearized with SmaI. To obtain pMB2007, pMB2006 was linearized by PCR with primers 8-13/8-14 (8878 bp) and subsequently fused to the *tetA* gene fragment (1349 bp; primers 8-15/8-16) by isothermal assembly.

**Construction of plasmids expressing NifL-NifA from other Proteobacteria.** A plasmid, pPR34, for the expression of the *nifLA* operon from *A. vinelandii* has been previously constructed (13) allowing activation of a *nifH::lacZ* fusion in *E. coli* ET8000. To compare the activity of NifL-NifA proteins from *A. olearius* DQS4 (Ao) and *P. stutzeri* A1501 (Ps) with the archetypal proteins from *A. vinelandii* (Av), while avoiding differences that may arise from plasmid copy number or protein expression levels, we constructed a series of plasmids derived from pPR34 to express *A. olearius* NifL-NifA and *P. stutzeri* NifL-NifA. The pPR34 backbone, excluding the *nifLA* genes from *A. vinelandii,* was amplified with primers 3-75 and 4-14 to generate a 2421 bp fragment. To allow isothermal enzymatic assembly, 15-25 bp overlaps to the linearized pPR34 backbone were introduced into the 5' and 3' ends of *nifL* and *nifA,* respectively, from *A. olearius* or *P.stutzeri*. The derived plasmids, pMB1806 (*A. olearius* NifL-NifA) and pMB1804 (*P. stutzeri* NifL-NifA) were the templates for the introduction of point mutations by overlapping PCR mutagenesis to generate plasmids pMB1807 (*A. olearius nifA-E351K*) and pMB1805 (*P. stutzeri nifA E356K*). All plasmids and primers are listed in Tables S2 and S3, respectively.

**Recombinant DNA work.** General molecular biology techniques were performed according to established protocols (1). Enzymatic isothermal assembly (14) was performed with the NEBuilder® HiFi DNA Assembly Master Mix (NEB #E2621). Site-direct mutagenesis by overlapping PCR was performed as described previously (15). High-fidelity DNA polymerase and restriction enzymes were provided by New England Biolabs. DNA purification was performed using commercially available kits provided by Macherey-Nagel. Sanger DNA sequencing and oligonucleotide synthesis was conducted by Eurofins MWG Operon.

**Quantification of ammonia.** Ammonia from culture supernatants was quantified by the indophenol-blue method modified from (16, 17). In brief, 750 µL of sample (usually 50 µL of supernatant diluted to 750 µL in $H_2O$), or the calibration curve standards, were mixed with 150 μL of sodium phenate (0.25 M phenol, 0.3 M NaOH), 225 μL of 0.66 mM sodium nitroprusside dihydrate and 225 μL of 1-1.5% sodium hypochlorite, in this order. The final composition of the reaction mix was approximately: 28 mM phenol, 33 mM NaOH, 25 mM sodium hypochlorite and 0.11 mM sodium nitroprusside. Samples were incubated for approximately 40 minutes at room temperature and the resulting indophenol-blue quantified at 625 nm. The calibration curve used known concentrations of $NH_4Cl$ ranging from 0 to 0.3 mM (in 0.05 increments) from a 1 mM standard.

**β-galactosidase activity.** β-galactosidase activity assays were performed as described previously (18) and reported in Miller Units for *E. coli* or as specific activity units (nmol ONP.mg protein$^{-1}$.min$^{-1}$) for *A. vinelandii*. Growth conditions for assays in *E. coli* ET8000 were as previously established (4, 13, 19), except that 6 mL screw capped bijou universals were filled to the brim to enable anaerobic conditions (20). For assays involving *A. vinelandii* strains, cultures were prepared as for the nitrogenase activity measurements as described below. Specific β-galactosidase activity was calculated using an ONP (o-nitrophenol – Sigma# N19702) calibration curve prepared in the reaction buffer containing the same amount of $Na_2CO_3$ used to stop β-galactosidase development, as addition of $Na_2CO_3$ intensifies ONP colour development (18). Whole cell protein concentration was determined by the Bradford method (21) after overnight cell lysis in 0.1 mM NaOH.

**Nitrogenase activity.** *In vivo* nitrogenase activity in *A. vinelandii* and *P. stutzeri* was measured by the acetylene reduction assay (22, 23) in cultures growing in NIL liquid media as indicated below. After incubation of cells with acetylene for 0.5- to 1-hour (*A. vinelandii*) or 4-18 hours (*P. stutzeri*), ethylene ($C_2H_4$) was quantified using a Perkin Elmer Clarus 480 gas chromatograph equipped with a HayeSep® N (80-100 MESH) column. The injector and oven temperatures were kept at 100 °C, while the FID detector was set at 150 °C. The carrier gas (nitrogen) flow was set at 8 - 10 mL/min.

Nitrogenase activity is reported as nmol of $C_2H_4$. mg protein$^{-1}$.min$^{-1}$. The ethylene calibration curve was prepared from chemical decomposition of ethephon (Sigma #C0143) in a 10 mM $Na_2HPO_4$ pH = 10.7 as described previously (24). As *A. vinelandii* is able to fix nitrogen in air (21 % $O_2$) 20 mL cultures were grown in NIL media as above in 100 mL conical flasks in an open atmosphere at 250 rpm until the desired $O.D_{600nm}$ was reached. Immediately before the acetylene reduction assay, the flask was stoppered with rubber septa (Suba-Seal® nº37), 10% acetylene injected, and the ethylene formed analysed after 0.5- to 1-hour of incubation. Given the microaerobic lifestyle of *P. stutzeri*, the flasks were prepared at defined initial oxygen concentrations of 4% or 8% to provide appropriate conditions for rapid nitrogenase de-repression and ammonia excretion, respectively. 50 mL of cells pre-cultured in LB overnight were collected by centrifugation and resuspended in 10-15 mL of UMS basal medium without carbon source to wash away excess nitrogen. 20 mL of bacterial suspension at an initial $O.D_{600nm}$ of 0.2-0.4 (aiming for a final $O.D_{600nm}$ of 0.55-0.6 at the end of the incubation period) was prepared in UMS-PS medium, transferred to 100 mL conical flasks and stoppered with the rubber septa. The flasks were flushed with nitrogen for 20-25 min prior to adjusting the oxygen concentration by injecting back a defined amount of air into the flask. After oxygen concentration adjustment, the flasks were incubated for 4 hours at 120 rpm and 30ºC and then 10% acetylene was injected. The ethylene formed was analysed after 4-18 hours.

**Glutamine synthetase activity.** Glutamine synthetase biosynthetic (GSB) and transferase (GST) activities were determined using previously described protocols (25, 26) with minor modifications. Prior to the assay, 20 mL of cells grown under the conditions described earlier, were quenched by addition of 2 mL of CTAB (1 mg/mL) under constant agitation (250 rpm) for 3 minutes. Cells were then immediately collected by centrifugation at 4°C, washed once in 20 mL of 1% KCl, collected by centrifugation once more, and finally resuspended to a final volume of 1 mL in 1% KCl. The resulting cell extract was kept on ice, and immediately used for the activity assays.

For the biosynthetic activity assay (GSB), 40 µL of cell extract was added to 400 µL of GSB assay mix (234 mM imidazole hydrochloride pH 7.4, 58.6 mM hydroxylamine hydrochloride, 70.4 mM magnesium chloride hexahydrate, 209 mM L-sodium glutamate and 117.3 µg/mL of CTAB) and equilibrated at 37°C for 5 minutes. The reaction was started by the addition of 60 µL 0.2 M ATP. For the transferase activity assay (GST), 50 µL of cell extract was added to 400 µL of GST assay mix (168.6 mM imidazole hydrochloride pH 7.15, 22.2 mM hydroxylamine hydrochloride, 0.34 mM manganese chloride, 31.5 mM sodium arsenate pH 7.15, 0.44 mM ADP and 111 µg/mL of CTAB) and equilibrated at 37°C for 5 minutes. The reaction was started by the addition of 50 µL 0.2 M L-glutamine. Both reactions were incubated at 37°C for 30-40 minutes and stopped by the addition of 1 mL of the stop mix (55 g/L of $FeCl_3.6H_2O$, 20 g/L of trichloroacetic acid, 21 mL/L of concentrated HCl). The product of both reactions, L-Glutamyl-γ-Hydroxamate (LGH), was quantified at 540 nm. The activity is reported as nmol LGH.mg $protein^{-1}.min^{-1}$.

**Fluorometric quantification of 2-oxoglutarate.** Prior to metabolite extraction, 40 mL of cells were rapidly vacuum filtered through a 0.22 µM cellulose acetate filter coupled to a 50 mL falcon tube (Corning # CLS430320).  The retained cell biomass was recovered from the filter in 2 mL of 0.3 M $HClO_4$. After recovery, 1.7 mL of the suspension was rapidly transferred to a chilled 2 mL tube, vigorously vortexed and centrifuged (17000 x g, 4ºC, 5 min) to remove cell debris. Then, 1.5 mL of the supernatant was transferred to a new tube and neutralized with 230 µL of 2M $K_2CO_3$ followed by a 5 min incubation on ice. To precipitate the excess $KClO_4$ formed, the extract was centrifuged (17000 x g, 4°C, 5 min) and the supernatant carefully transferred to a fresh tube avoiding touching the white precipitate ($KClO_4$). 1-2 µL of the neutralized extract was used to estimate the pH (7.5 – 8.5) using pH strips. The volume of 2M $K_2CO_3$ used was optimized to achieve complete neutralization. The extracts were then stored at -80°C or used immediately for fluorometric quantification of 2-oxoglutrate as described previously (27–29). In brief, the detection assay was performed in 300 µL final volume and contained 100 mM imidazole-acetate buffer pH 7.0, 60 mM ammonium acetate, 10 µM NADH, 100 µM ADP and 0.075 µg of glutamate dehydrogenase (Sigma #G7882-100MG). Reactions were started by adding the enzyme and were incubated at 25°C for 20 minutes or until stabilization of the NADH fluorescence decay. The NADH fluorescence decay calibration curve was interpolated with 2-oxoglutarate standards ranging from 0.18 to 2.7 nmol prepared in a "mock" extraction solution obtained from $HClO_4$ neutralization with $K_2CO_3$ in the same fashion as in the metabolite extraction procedure above. 200 µL (out of 300 µL) of the reactions were used to measure the NADH decay ($\lambda_{ex}$: 340nm and $\lambda_{em}$: 460nm) using a 96-well black plate with clear bottom (Corning # CLS3603-48EA) and the BMG CLARIOstar plate reader. The internal metabolite concentration was calculated assuming that the cell-bound water content is approximately four times the bacterial dry weight (30).

**RNA purification and quantitative RT-PCR.** Prior to RNA purification, *A. vinelandii* DJ (wild type) and the *nifA-E356K* mutant (EK) were submitted to an ammonium switch as described previously (31). Briefly, cells growing under ammonium excess were collected by centrifugation and resuspended to an $O.D_{600\ nm}$ of 0.5 in fresh medium without added ammonium and subsequently incubated for 4 hours prior to RNA extraction. For *P. stutzeri*, the RNA was purified from cultures prepared exactly as described for the nitrogenase activity assay above. To ensure preservation of intracellular RNA, the cultures were immediately mixed with 1/5 of stop solution (5% Phenol saturated with 0.1 M citrate pH 4.3, 95% ethanol) (32) and then rapidly chilled on ice for 20 minutes. RNA was purified using the TRI Reagent® (Sigma #T9424) following manufacturer instructions. Genomic DNA was removed using TURBO DNA-*free*™ DNAse (Ambion #AM1907) following the rigorous DNase treatment according to the manufacturer. cDNA synthesis was performed with SuperScript™ II Reverse Transcriptase (Invitrogen #18064014) using 0.1-1 µg of total RNA as recommended by the manufacturer. The resulting cDNA was diluted 5 to 20-fold (to fit the genomic DNA calibration curve) and 2 µL used as template in a 20 µL qPCR performed with the SensiFAST™ SYBR® No-ROX Kit (#BIO-98005) reagent and the Bio-Rad CFX96 instrument. Absolute quantification of target genes (*nifH, nifL* and *nifA*) alongside the normalizing house-keeping gene (*gyrB*) was performed according to (33) using a $C_q$ calibration curve interpolated using serial dilutions of purified genomic DNA from *A. vinelandii* and *P. stutzeri*. Relative quantification was performed by the $2^{\wedge-\Delta Cq \Delta Cq}$ method according to (34). Primers were designed and validated according to (35) ensuring comparable efficiencies and specificity as judged by the presence of a single peak in the melting curve. The primers used are listed in Table S3.

### Supplementary Figures

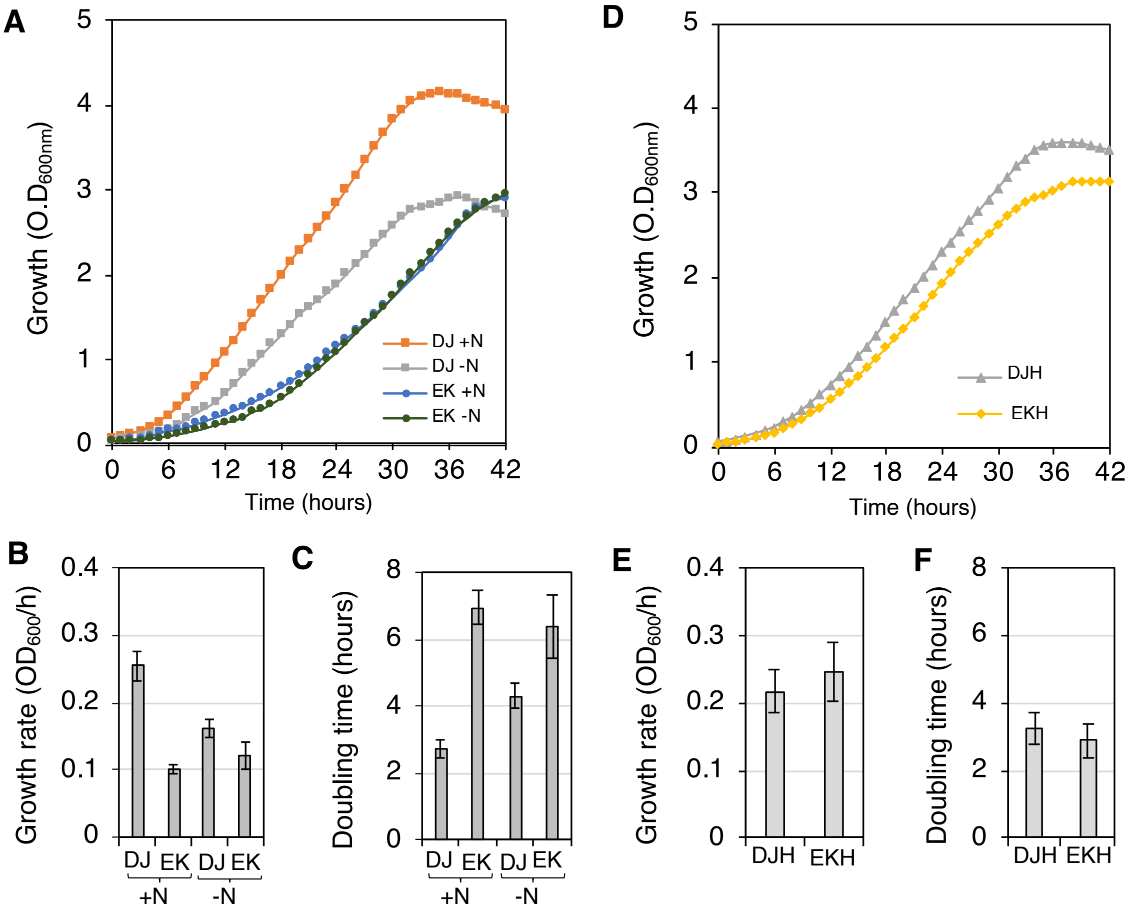


**Fig. S1 – The *A. vinelandii nifAE356K* strain (EK) has a growth penalty that is dependent on overexpression of the nitrogenase structural genes (*nifHDK*).** (A) Strains were grown in MBB media supplemented with 2% sucrose either in the presence (+N) or absence (-N) of 25 mM ammonium acetate. The growth rates (B) and doubling times (C) in the exponential phase of growth are also shown. (D) Strains carrying a *nifH* insertion were grown only in the presence of 25 mM ammonium acetate, given that they are unable to grow diazotrophically. The growth rates (E) and doubling times (F) in the exponential phase of growth were calculated from the data in (D). Cells were assayed for growth on a 24-well microplate (Greiner-Bio one #662160) using the Biotek EON plate reader as described in the Materials and Methods section. The absorbances recorded at 600 nm were corrected to a pathlength of 1 cm.

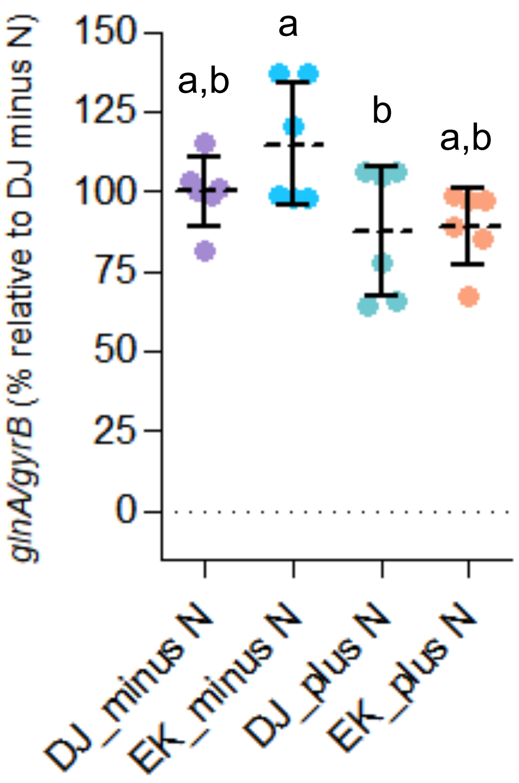


**Fig S2 – Levels of *glnA* transcripts in *A. vinelandii* strains grown under two nitrogen regimes.** The strains were grown in minimal media supplemented with 2% sucrose under diazotrophic conditions (-N) or in the presence of excess ammonium chloride (+N). Data is relative to the maximum level of detected transcripts in the wild type under diazotrophic conditions (DJ -N) estimated from absolute quantification. The data is representative from 2 independent RNA purifications performed in technical triplicate. Plots followed by different letters are statistically different according to ANOVA with post-hoc Tukey's HSD.

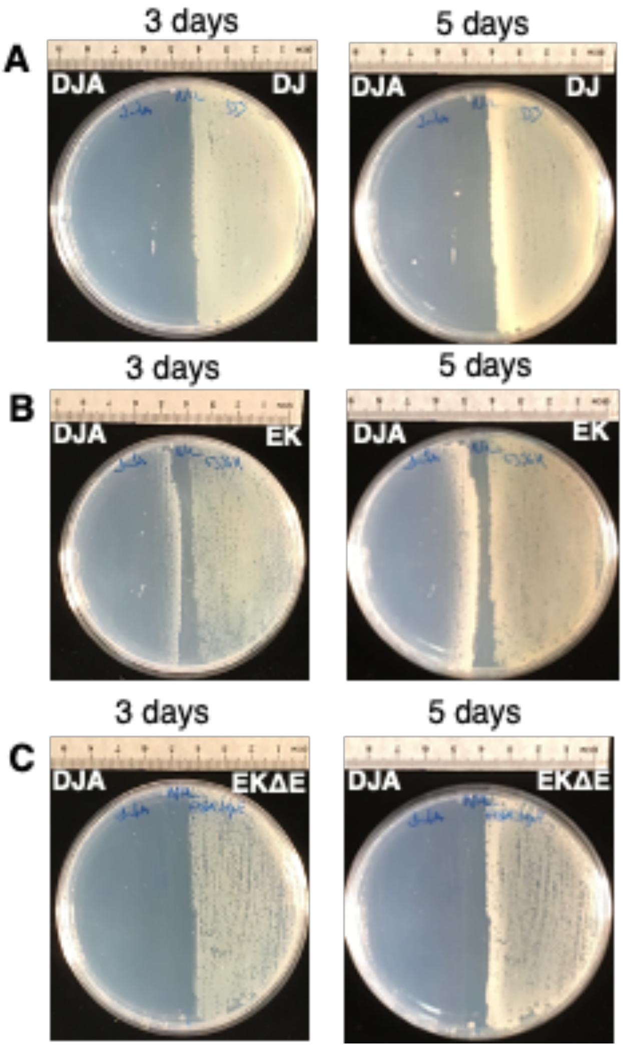


**Fig. S3 – Ammonia excretion ceases in a strain unable to adenylylate glutamine synthetase.** Cells were spread on opposite sides of an agar plate containing NIL media with 2% sucrose but without fixed nitrogen. The *nifA* deletion (DJA) is unable to grow unless a source of fixed nitrogen is provided. When spread opposite to the wild type strain (DJ) the *nifA* deletion strain (DJA) was unable to grow (A). In contrast, DJA grew when spread opposite to the strain EK due to the diffusion of excreted ammonium (B). When the *glnE* gene is deleted in the *nifAE356K* background (strain EKΔE), ammonium excretion is impaired and so is growth of the *nifA* deletion strain (C).

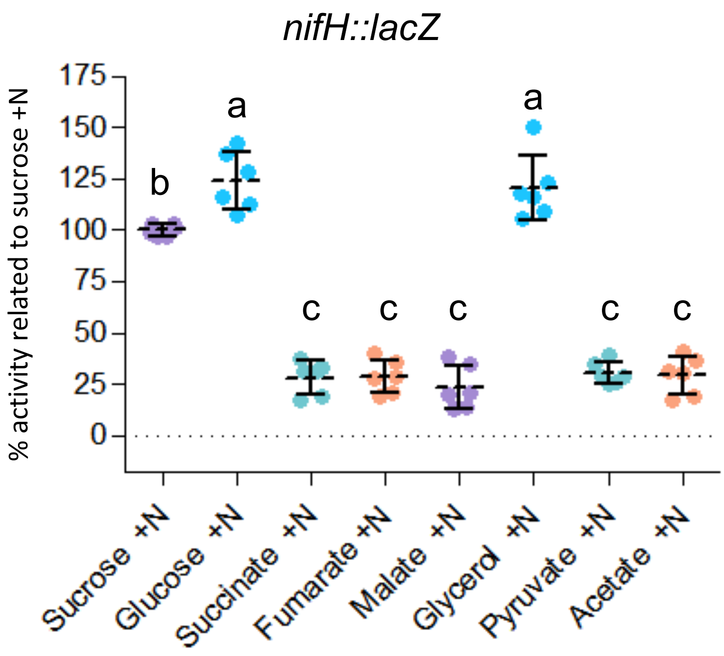


**Fig. S4 – Activation of *nif* gene expression by NifA-E356K in different carbon sources as reported by a *nifH::lacZ* fusion in *A. vinelandii* (strain** EKHZ**).** Cell suspensions (O.$D_{600}$ = 0.1) were spotted as 10 µL drops (triplicate) on minimal solid media plates supplemented with 10 mM $NH_4Cl$ and the carbon sources indicated. After 18-36 hours incubation at 30ºC, the grown bacterial biomass from triplicate spots were pooled and resuspended in 1 mL of PBS buffer. The β-galactosidase activity was performed using 100 µL of the PBS resuspended cells. Data is relative to the maximum level of detected activity in sucrose, calculated from specific β-galactosidase activity (1287.08 ±242.76 nmol ONP. mg protein$^{-1}$. min$^{-1}$). The data is representative from 3 independent experiments performed in technical duplicate. Plots followed by different letters are statistically different according to ANOVA with post-hoc Tukey's HSD.

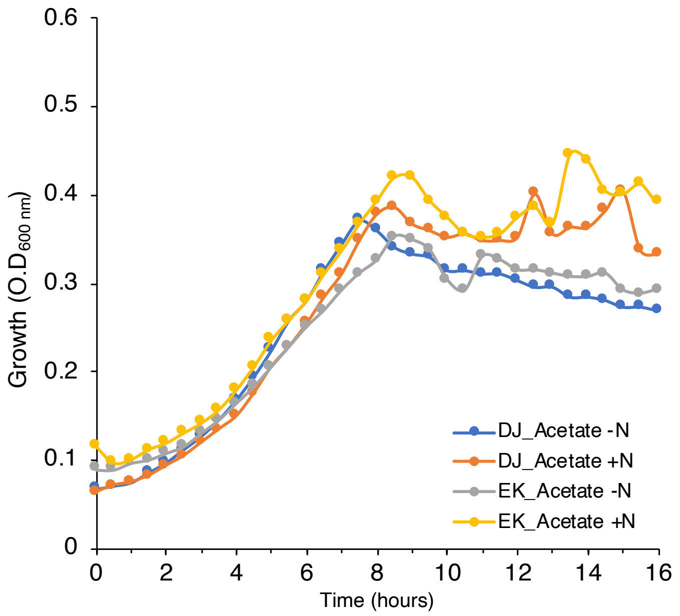


**Fig. S5 – Growth profile of the *A. vinelandii nifA-E356K* (EK) strain compared to the wild type (DJ) using acetate as a carbon source.** Cells were grown in MBB media supplemented with 30 mM acetate as carbon source without added ammonium (-N) or with 10 mM ammonium chloride (+N). Growth was assayed as described in Fig. S1.

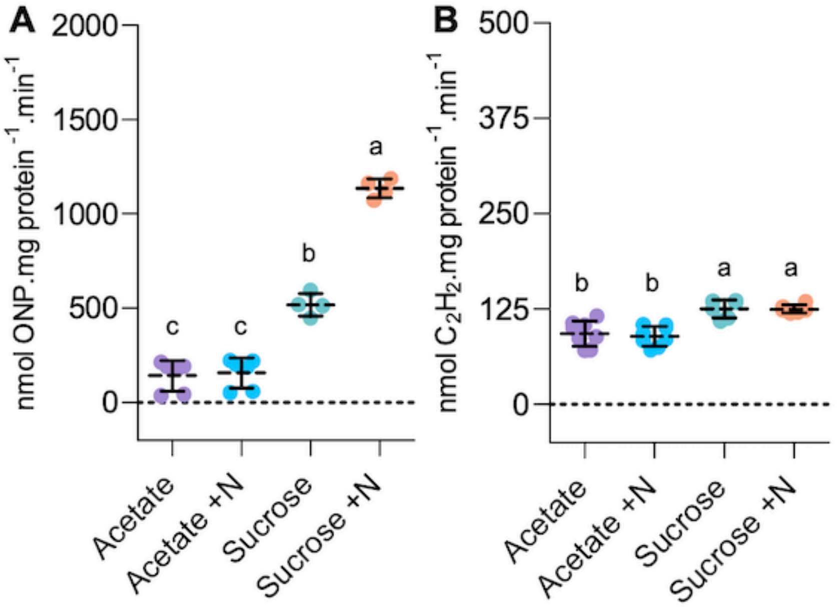


**Fig. S6 – Profile of nitrogenase expression and activity in a *nifL*::Kan[r] insertion strain in different carbon and nitrogen regimes.** The *nifL* disrupted strain (AZBB163) was modified to encode a *nifH::lacZ* fusion (strain 163HZ) allowing ready comparison of nitrogenase expression in (A) and nitrogenase activity in (B). Plots followed by different letters are statistically different according to ANOVA with post-hoc Tukey's HSD.

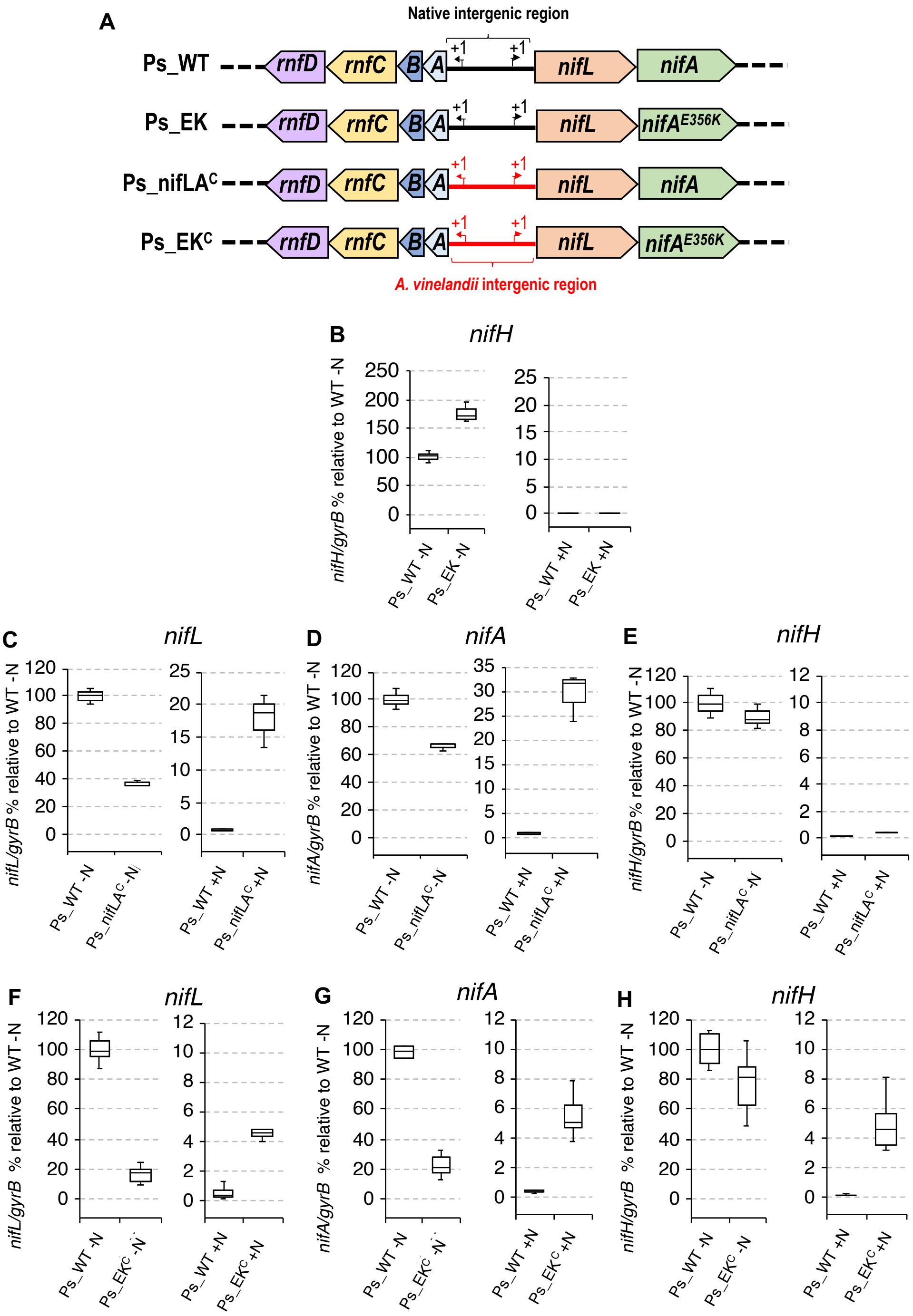


**Fig. S7 – Levels of *nif* gene transcripts in various *P. stutzeri* strains.** (A) Diagram depicting the genotypes of the *P. stutzeri* strains analysed. Drawings are not to scale. (B) *nifH* transcripts in the strain Ps_EK compared to the wild type (Ps_WT). (C-E) levels of *nifL*, *nifA* and *nifH* transcripts in the strain Ps_nifLA$^C$ compared to Ps_WT. (F-H) levels of *nifL*, *nifA* and *nifH* transcripts in the strain Ps_EK$^C$ compared to Ps_WT. Strains were grown on 2% glucose under diazotrophic conditions (-N) or in the presence of excess fixed nitrogen (5 mM $NH_4Cl$, +N). In each case the data is relative to the maximum level of detected transcripts under fully derepressing conditions (Ps_WT -N) estimated from absolute quantification.

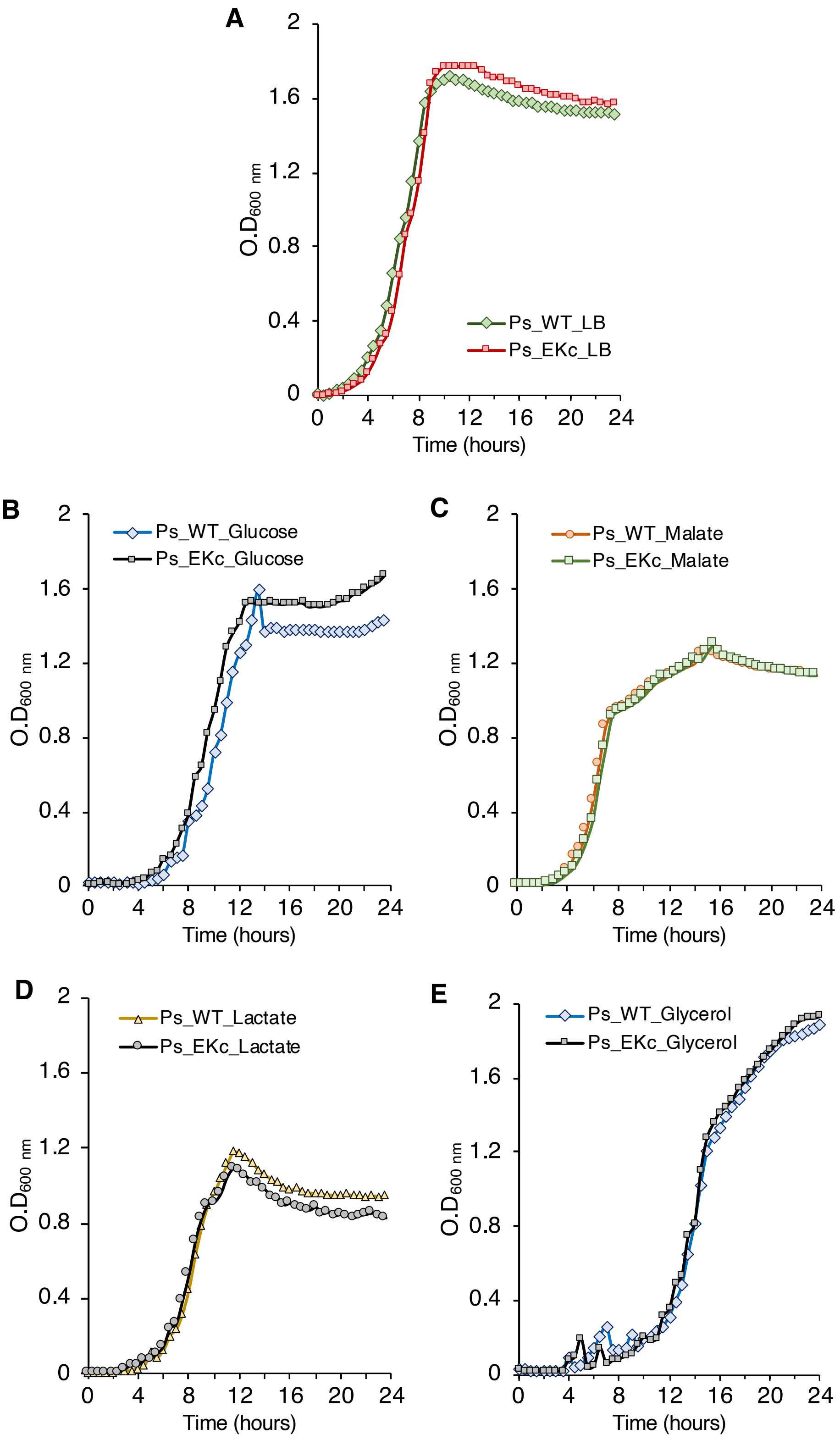


**Fig. S8 – Growth profiles of *P. stutzeri* A1501 (wild type, Ps_WT) and the *nifA-E356K* mutant (Ps_EK$^{C}$) under nitrogen replete conditions.** Growth as assayed in LB media (A) or in UMS-PS medium supplemented with 30 mM glucose (B), 45 mM malate (C), 60 mM lactate (D) or in 60 mM glycerol (D). In (B-D) the nitrogen source used was 5 mM $NH_4Cl$.

### Supplementary Tables

Table S1 - Strains used in this study

| Strain | Relevant characteristic | Source |
|---|---|---|
| *A. vinelandii* | | |
| DJ | ATCC BAA-1303. High frequency transforming variant of *A. vinelandii* UW generated in 1984 by Dennis Dean | (11) |
| DJA | DJ derived. 1401 bp deletion of *nifA* gene. Mo-Nif-, $Rif^R$ | R.D. lab |
| DJH | DJ derived. *nifH::tetA*. $Mo\text{-}Nif^-$, $Tc^R$ | This study |
| DJHZ | DJ derived. *nifH::lacZ* inserted into the *algU* locus. $Tp^R$ | This study |
| EK | DJA derived. *nifA* deletion rescued by insertion of *nifA-E356K* | This study |
| EKH | EK derived. *ΔnifH::tetA* insertion into the native locus. $Tc^R$ | This study |
| EKHZ | EK derived. *nifH::lacZ* fusion inserted into the *algU* locus. $Tp^R$ | This study |
| EKΔE | EK derived. *ΔglnE::tmp* insertion into the native locus. $Tp^R$ | This study |
| AZBB163 | *nifL::Km*$^R$. Spontaneous mutant from AZBB150 resulting in the $Nif^+$ phenotype | (36) |
| 163HZ | AZBB163 derived. *nifH::lacZ* inserted into the *algU* locus | This study |
| *P. stutzeri* | | |
| A1501 | CGMCC 0351, wild-type, isolated from rice roots inoculated with strain A15 | (37, 38) |
| Ps_*nifLA*$^C$ | A1501 derived. *rnf-nifLA* intergenic region swapped by the reciprocal *A. vinelandii* region. | This study |
| Ps_EK$^C$-tetA | Ps_*nifLA*$^C$ derived, but *Ps_nifAE356K* + *tetA* ($Tc^R$) | This study |
| Ps_EK$^C$ | Ps_EK$^C$-tetA derived, but *tetA* cured with pMB2006 | This study |
| Ps_EK | A1501 derived, Ps_*nifAE356K* | This study |
| *E. coli* | | |
| ET8000 | *rbs lacZ::*IS1 *gyrA hutC.* Used for β-galactosidase assays | (39) |
| NEB 5-alpha | Derivative of DH5α. T1 phage resistant and *endA* deficient. Used for cloning and plasmid maintenance | NEB |
| ST18 | *E. coli* S17λpir Δ*hemA.* Used for conjugations | (3) |

Abbreviations: $Mo\text{-}Nif^-$: no molybdenum-dependent nitrogenase activity; Rif: rifampicin, Km: kanamycin, Tc: tetracycline, Tp: trimethiorim, $^R$: resistance

Table S2 – Plasmids used in this study

| Plasmid | Relevant characteristic | Source |
|---|---|---|
| pAAS1544 | $Cb^R$. Derived from pPR54 (13) encoding *A. vinelandii* NifL (147-519) and NifA-E356K. | Adriano Stefanello unpublished |
| pPR34 | $Cb^R$. *A. vinelandii nifLA* translated from the natural ribosome binding site of *nifL* in pT7-7 | (13) |
| pPMA | $Cb^R$. Derived from pPR34, but encoding *nifA-E356K* | (19) |
| pRT22 | $Cm^R$. *pnifH::lacZ* in pACYC184 | (40) |
| pBlueScript SK II + | $Cb^R$. Cloning vector | (41) |
| pUC19 | $Cb^R$. Cloning vector | (42) |
| pK18mobsacB Km | $Km^R$, Mob. Suicide vector for gene replacement. sacB gene for counter selection | (43) |
| pALMAR3 | $Tc^R$. Source of tetracycline resistance gene (*tetA*) | (44) |
| pUC18T-mini-Tn7T-Tp | $Tp^R$. Source of the trimethoprim resistance gene (*tmp*) | (45) |
| pMB1724 | $Cb^R$. 1893 bp PCR fragment corresponding to A. vinelandii *nifHD* region cloned into BamHI/HindIII sites of pBlueScript II + | This study |
| pMB1725 | $Cb^R$. $Tc^R$. *tetA* gene amplified by PCR from pALMAR3 inserted into BglII and EcoRI sites of pMB1724 | This study |
| pMB1804 | $Cb^R$. Derived from pPR34 encoding *P. stutzeri* A1501 *nifL-nifA* | This study |
| pMB1805 | $Cb^R$. Derived from pPR34 encoding *P. stutzeri* A1501 *nifL-nifA-E356K* | This study |
| pMB1806 | $Cb^R$. Derived from pPR34 encoding *A. olearius* DQS-4 *nifL-nifA* | This study |
| pMB1807 | $Cb^R$. Derived from pPR34 encoding *A. olearius* DQS-4 *nifL-nifA-E351K* | This study |
| pMB1816 | $Cb^R$, $Tp^R$. *A. vinelandii nifH::lacZ* fusion flanked by homology regions for integration into *algU* genome locus | This study |
| pMB1840 | $Km^R$. $Tp^R$. *A. vinelandii glnE* deletion. Fragments upstream (906 bp) and downstream (752 bp) of *glnE* were fused to the *tmp* gene, and inserted into pk18mobsacBKm cut with SmaI | This study |
| pMB2005 | $Km^R$. The *rnf-nifLA* intergenic region (444 bp) from *A. vinelandii* was fused downstream to a fragment of the *P. stutzeri rnfAB* genes (1160 bp) and upstream to a fragment of the *P. stutzeri nifL* gene (1638 bp) and inserted into pk18mobsacBKm cut with SmaI | This study |
| pMB2006 | $Km^R$. Ps-*nifLAE356K* fragment (2364 bp) from pMB1805 was fused to an 847 bp fragment downstream Ps-*nifA* and inserted into pk18mobsacBKm cut with SmaI. Construct to recover *tetA* from Ps_EK$^C$-*tetA* to generate Ps_EK$^C$ | This study |
| pMB2007 | $Km^R$, $Tc^R$. The plasmid pMB2006 was linearized by PCR and fused to a fragment encoding the *tetA* gene (1349 bp). Construct to generate Ps_EK$^C$-*tetA* | This study |

Abbreviations: Cb: carbenicillin, Cm: chloramphenicol, Km: kanamycin, Tc: tetracycline, Tp: trimethiorim, $^R$: resistance.

Table S3 - Primers used in this study

| ID | Sequence 5' -> 3' | Application |
|---|---|---|
| AAS-3 | **GAATGCCCATGAATGCAACCAT** | Amplification of Av-*nifA* for genotyping |
| AAS-55 | **TCAGATCTTGCGCATGTGGATGT** | |
| 2-43 | **AAAAGGATCCCTCATGTTTGACAGCTTATC** | Amplifies the *tetA* cassette from pALMAR3 for Av-*nifH* insertion |
| 2-44 | **AAAAGAATTCTTGATTGGCTCCAATTCTTG** | |
| 2-45 | **AAAAGGATCCGTCACTGAACTCTCCTGCTGAGG** | Amplifies 1874 pb of Av-*nifHD* (position 136301 – 138174) |
| 2-46 | **AAATAAGCTTTCCACTTCGTCGATCAGTTTGGCG** | |
| 5-49 | **GAGTTGTTCGCTTTCTACGGAATCATTGGTGATTTCGG** | Amplifies 409 bp corresponding to the Av-*nifH* promoter (position 136401 – 136809) |
| 5-50 | **TGTAAAACGACGGGATCCCCGGACTTACCGATACCACC** | |
| 5-46 | **GGGGATCCCGTCGTTTTACAACGTCGTG** | Amplifies the *lacZ* gene (3057 bp) and from pRT22 |
| 5-464 | **CTAGCGGTTATTATTATTTTTGACACCAGACCAACTG** | |
| 7-28 | **CGAATTCGAGCTCGGTACCCCCATGGAAACGGTGGGGAACTG** | Amplifies 906 bp upstream Av-*glnE* (position 4547284 – 4548189) |
| 7-29 | **AATTATCCATACTGTCGCCGAGCGTGCC** | |
| 7-30 | **CGGCGACAGTATGGATAATTCACGAACC** | Amplifies a 631 bp fragment from pUC18T-mini-Tn7T-Tp encoding to the *tmp* resistance gene |
| 7-31 | **CCGGCTGCTTTAATAACCGCTAGATAATTCTTAG** | |
| 7-32 | **GCGGTTATTAAAGCAGCCGGGCGTGGTG** | Amplifies 752 bp downstream Av-*glnE* (position 4543778 – 45444529) |
| 7-33 | **GTCGACTCTAGAGGATCCCCGCCTTCCGGATCGAGCAGC** | |
| 3-75 | **AAGCTTATCGATGATAAGCTG** | Amplification of 2421 bp from pR34 excluding Av-*nifLA* |
| 4-14 | **ATGGTGCCTCGTCTATCCGAA** | |
| 4-17 | **GGAAGCTCGCATGAACGCCACATTCGCC** | Amplification of Ps-nifA for constructing pMB1804 |
| 4-18 | **CGATAAGCTTTCAGATCTTGCGCATATGAATG** | |
| 4-19 | **GAGGCACCATATGGCTTTGCAACGGATACC** | Amplification of Ps-nifL for constructing pMB1804 |
| 4-20 | **TGGCGTTCATGCGAGCTTCCCCTGTCAG** | |
| 4-21 | **CGCGACCTGAAGCATGAGGTGGAG** | Introducing E356K mutation into Ps-nifA |
| 4-22 | **CGCGACCTGAAGCATGAGGTGGAG** | |
| 3-74 | **CAGCTTATCATCGATAAGCTTTCAGATCTGCCGCACCTTGAT** | Amplification of Ao-nifA for constructing pMB1806 |
| 3-74B | **GACCTGGAGGGCGGTCGATGAGCGCGGCCGGTCCGATG** | |
| 4-12 | **CATCGGACCGGCCGCGCTCATCGACCGCCCTCCAGGTC** | Amplification of Ao-nifL for constructing pMB1806 |
| 4-13 | **TTCGGATAGACGAGGCACCATATGGGCGCTGTCGCCGACG** | |
| 2-37 | **ACCAACCGCGACCTCAAGCTCGAGGTCGAAGCG** | Introducing E351K mutation into Ao-nifA |
| 2-38 | **CGCTTCGACCTCGAGCTTGAGGTCGCGGTTGGT** | |
| 8-1 | **TGATTACGAATTCGAGCTCGGTACCCTCAGTCTTCGGTTTCCGCCGCT** | Amplification of 847 bp downstream Ps-*nifA* |
| 8-2 | **GCGCAAGATCTGAACGACCCGCCCG** | |

| | | |
|---|---|---|
| 8-3 | TCGGGCGGGTCGTTCAGATCTTGCG | Amplification of Ps-*nifLAE356K* (2364 bp) fragment from pMB1805 |
| 8-11b | CTCTAGAGGATCCCCACCGGGCTGCGCCAGCAA | |
| 8-13 | ACATGAGCCCGGGTCGGGC | Linearizes the pMB2006 by PCR for *tetA* insertion |
| 8-14 | CCAATCAAAGTCGGGCGGGTCG | |
| 8-15 | GCCCGACTTTGATTGGCTCCAATTCTTGGAGTG | Amplifies the *tetA* cassette from pALMAR3 for insertion into pMB2006 |
| 8-16 | GACCCGGGCTCATGTTTGACAGCTTATCATCGATTAGC | |
| 8-17 | TCGAGCTCGGTACCCTCAGGCGGCCCT | Amplification of Ps-*rnfAB* (1160bp) |
| 8-18 | GAGGTCGCTTATGGAATATGCGCTGTTTCTGATCG | |
| 8-19 | CATATTCCATAAGCGACCTCACCTGCT | Amplification of the *A. vinelandii rnf-nifLA* intergenic region (444 bp) |
| 8-20 | AAAGCCATGCTGTGCCTCGTCTATCCA | |
| 8-21 | GCACAGCATGGCTTTGCAACGGATACCG | Amplification of Ps-*nifL* (1638 bp) |
| 8-22 | GACTCTAGAGGATCCCCTCAGCTGGCCGAGAAGGG | |
| M13F (-47) | CGCCAGGGTTTTCCCAGTCACGAC | Sequencing and PCR linearization of constructs |
| M13R (-48) | AGCGGATAACAATTTCACACAGGA | |
| Av-RTgyrBF | CAAGAAGCACAAGGTGACGA | RT-qPCR *A. vinelandii gyrB* |
| Av-RTgyrBR | TGGTCTGCGAACTGAACTTG | |
| AvRTnifH-3F | CAGCCCTGGCTGAGATGGG | RT-qPCR *A. vinelandii nifH* |
| AvRTnifH-3R | ATGGTGTTCTGGGCCTTGGA | |
| Av-RTnifLF | CTGCTGACCATCAACGACAT | RT-qPCR *A. vinelandii nifL* |
| Av-RTnifLR | ATGCCTTCCAGCAGCTCTT | |
| Av-RTnifAF | GCAAGTACGGCTTCGAGAAC | RT-qPCR *A. vinelandii nifA* |
| Av-RTnifAR | GTACGGTGCTGTTCCACTTG | |
| Ps-RTgyrB2F | AAACCATCCGCCGAGACCTT | RT-qPCR *P. stutzeri gyrB* |
| Ps-RTgyrB2R | CCCACCCCGGAGTTGAGAAA | |
| Ps-RTnifH1F | CCACGACCCAGAACCTCGTG | RT-qPCR *P. stutzeri nifH* |
| Ps-RTnifH1R | GAGTGCAGGATCAGGCGAGT | |
| Ps-RTnifL2F | AGGACATGCACATGACCCAGG | RT-qPCR *P. stutzeri nifL* |
| Ps-RTnifL2R | CACGGCTTGCTGGAACACTT | |
| Ps-RTnifA7F | TCCTCAAGCATGGCAACAGC | RT-qPCR *P. stutzeri nifA* |
| Ps-RTnifA7R | GAAGGGCAGGTCCATGTCGTA | |
